# Resilience and vulnerability of neural speech tracking in children with cochlear implants

**DOI:** 10.1101/2024.02.22.581545

**Authors:** Alessandra Federici, Marta Fantoni, Francesco Pavani, Giacomo Handjaras, Evgenia Bednaya, Alice Martinelli, Martina Berto, Emiliano Ricciardi, Elena Nava, Eva Orzan, Benedetta Bianchi, Davide Bottari

## Abstract

Infants are born with biological biases that favour language acquisition. One is the auditory system’s ability to track the envelope of continuous speech, a pivotal feature for spoken language comprehension in adulthood. However, the extent to which neural speech tracking relies on postnatal auditory experience remains unknown. In this case-control study, we tested children with or without access to functional hearing in the first year of life after they received cochlear implants (CIs) for hearing restoration. We measured neural speech tracking in CI users with a congenital bilateral profound deafness (CD) or who acquired it later in development (AD; minimum auditory experience after birth 12 months), as well as in two groups of hearing controls listening to original (HC) or vocoded-speech (HC-v). Remarkably, neural speech tracking in children with CIs was unaffected by the absence of perinatal auditory experience. Regardless of deafness onset, CI users and HC exhibited a similar neural tracking magnitude at short timescales ∼50– 130 ms (P1_TRF_) of brain activity. However, this neural tracking phase (P1_TRF_) was delayed in CI users, and its timing depended on the age of hearing restoration. Conversely, at longer timescales ∼130–260 ms (N2_TRF_) of brain activity, speech tracking was substantially dampened in participants with CIs, thereby accounting for their comprehension deficits. Speech tracking in HC listening to vocoded-speech and in a phantom head-model with CIs suggested that neural processing differences between HC and CI children could not merely be explained by the degraded acoustic stimulation, nor by electrical artifacts of the implants. These findings highlight (i) the resilience of sensory components of neural speech tracking to the lack of hearing in the first year of life, (ii) the crucial role played when hearing restoration takes place in mitigating the impact of atypical auditory experience, (iii) the vulnerability of higher hierarchical levels of speech processing in CI users.

## Introduction

In the case of typical development, the auditory cortex tracks several speech sound features, such as signal amplitude modulations. These fluctuations, or speech envelope, have energy peaks around the syllabic rate and are a pivotal feature for speech comprehension. Adults can understand heavily degraded speech, provided the envelope is preserved^1^. On the contrary, however, its suppression impairs comprehension^2,3^. The primary role of this form of neural speech tracking is substantiated by studies revealing its presence in newborns and young infants^4–7^, thus suggesting a strong biological predisposition. Nonetheless, recent evidence suggests that linguistic experience in the first year of life modulates neural speech tracking^6^. Indeed, typical brain development requires temporal overlapping between neural system readiness and appropriate environmental statistics^8–10^. Yet, it is still unknown to what extent the development of neural speech tracking relies on postnatal hearing experience.

Individuals facing a period of sensory deprivation provide a unique opportunity for causally assessing whether neural functions have sensitive phases in which specific sensory input must be provided for shaping the associated neural circuitries^11^ or whether their development is mainly guided by biological predispositions instead. Following a period of profound bilateral sensorineural hearing loss (profound deafness from here onward), in which sounds cannot reach the auditory system, the cochlear implant (CI) provides the possibility of partial auditory restoration^12–14^. Cochlear implants rely substantially upon the sound envelope to convert continuous speech into electric impulses^15^. However, despite efficiently conveying this information to the brain, speech comprehension outcomes are very heterogeneous^16^. Crucial for understanding language acquisition variability is the age at which hearing is restored with cochlear implantation; results suggested the sooner, the better^17–20^. Seminal studies uncovered the fact that the latency of auditory responses in individuals using CIs falls within typical developmental trajectory only when children are implanted before 3.5 years of age^21–25^. However, the activity of the auditory cortex in CI individuals has been measured in reaction to simple and short-lived sounds, such as syllables. Thus, two key questions remain unanswered: (i) To what extent does the CI provide the possibility to develop hearing-like neural tracking of continuous speech? (ii) Is there a perinatal sensitive period during which auditory input is essential for its development?

To fill these gaps, we measured the degree of synchronization between brain activity and continuous speech (fitted and validated at the single participant level) in hearing children (HC), listening to either original or vocoded-speech (HC-v), and CI children with different onsets of bilateral profound deafness (congenital or acquired). The comparison between the neural speech tracking of children who were born with congenital deafness (CD) and experienced auditory deprivation during the first year of life, with children who acquired deafness (AD), born with some degree of functional hearing and whose profound deafness emerged only after one year of age, allowed to assess the role of the perinatal auditory experience (see Figure 1). Moreover, to control for the effect of degraded speech provided by the CI, speech neural tracking of CD and AD children was also compared with a group of hearing children who listened to vocoded-speech (HC-v). Participants’ age range (3–18 years old) was chosen to measure developmental trajectories of neural speech tracking in hearing and CI children.

**Figure 1.**
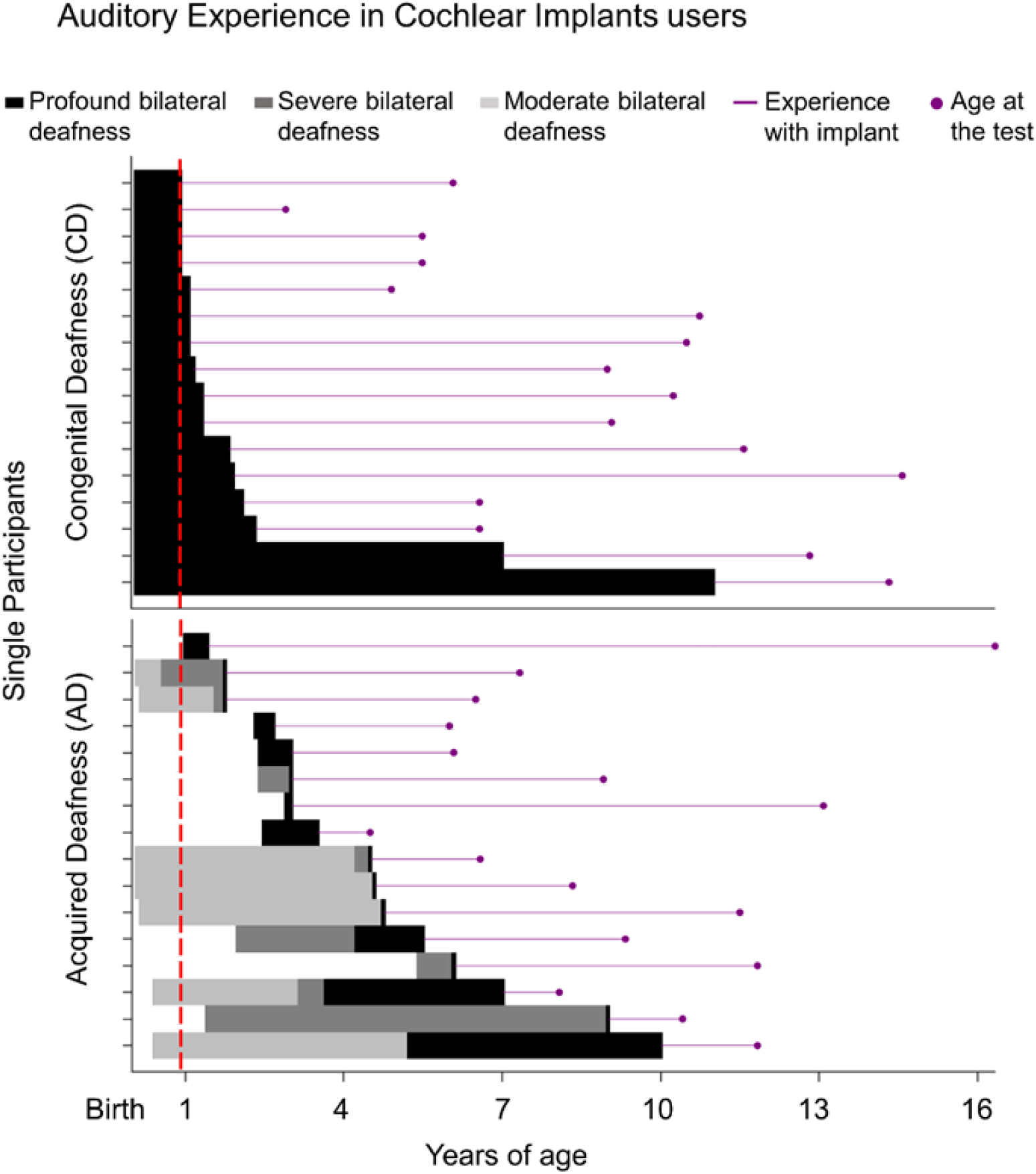
Auditory experience in CI users. The plot graphically represents the auditory experience that characterizes each CI participant. Periods of profound bilateral deafness are depicted in black, whereas severe and moderate bilateral deafness are rendered with different shades of gray. The red dashed line highlights the different hearing experiences between congenital deafness (CD) and acquired deafness (AD) groups: only the CD group faced auditory deprivation throughout the first year of life. The magenta lines illustrate experiences with implants until the day of testing, represented by the dot.

## Materials and methods

### Participants

A total of 97 children participated in the study. They were categorized according to their hearing status: cochlear implanted or hearing control children. A sample size of sixteen participants was estimated to reliably measure neural speech tracking through simulation using the data of ten hearing participants (see Supplementary Materials 1.1.1).

All CI children received cochlear implantation at least six months before the experimental test to ensure a consolidated auditory experience^21^. A total of forty-four children with cochlear implants were recruited at the Meyer Hospital of Florence (Italy) and the IRCCS Materno Infantile Burlo Garofolo of Trieste (Italy). The final sample of CI participants comprised thirty-two children (see Figure 1; Supplementary Materials 1.1.2 and 1.1.3 for the data exclusion and criteria): sixteen CD children (mean age=8.81 years; SD=3.52, eight females) and sixteen AD children (mean age=9.17 years; SD=3.15, nine females). As expected, the age of the diagnosis of profound bilateral deafness differed significantly between the two groups (t_(15.02)_=-7.35, p<0.001, d=-2.53, CI95=-3.58–-1.53). Importantly, no difference emerged between CD and AD in their experience with the implant (t_(30)_=1.53; p=0.136). The median age at cochlear implantation was 15 months (IQR=12, range: 11– 132) for the CD group and 48 months (IQR=36, range: 17–120) for the AD group (see Table S1 for detailed CI participants’ information).

A group of age- and gender-matched hearing children was recruited as the control group (HC, N=37; mean age=9.04 years; SD=4.10, seventeen females). Moreover, a group of sixteen hearing children was recruited to listen to vocoded-speech stimuli (HC-v, mean age=8.36 years; SD=1.49, six females). No significant difference emerged between the four groups, neither in age (F_(3,81)_=0.185; p=0.906) nor in gender (χ^2^_(3)_=1.203; p=0.752). HC children were recruited in Lucca and Milan (Italy). None of the children included in the final sample had any additional sensory deficits or neurological disorders (medical records and/or family reports). All participants were oralists; their first language (L1) was Italian (one CD, two AD, three HC, and one HC-v participants were bilingual).

The study was approved by the local Ethical Committee (see Supplementary Materials 1.1.4). Before participating in the experiment, written informed consent was signed by the participants’ parents and by the children themselves if they were older than seven years of age. The experimental protocol adhered to the principles of the Declaration of Helsinki (2013).

### Speech stimuli

The speech stimuli were 3-minute length stories read by a native Italian speaker. We chose different stories according to the children’s age to provide each participant with appropriate speech materials. Three different age ranges, 3–6, 7–10, and 11–15 years old, were defined according to Italian school cycles. For each age group, we selected ten stories from popular Italian books suitable for that age range (see Supplementary Materials 1.2.1 for more information about stimuli, their registration, preprocessing, and presentation).

Speech sound was delivered by a single front-facing loudspeaker (Bose Companion® Series III multimedia speaker system, country, USA) placed in front of the participants, approximately 70 cm distant from their heads. Loudness was delivered at ∼80 dB (Meterk MK09 Sound Level Meter).

We did not alter the sounds presented to the main hearing control group (HC). However, to control for the possible impact of receiving a degraded acoustic stimulation we also created a vocoded version of the stimuli for the HC-v group (see Supplementary Materials 1.2.2).

### Task and experimental procedure

Participants were asked to listen carefully to the stories while looking at a cross displayed at the center of a computer screen placed in front of them. At the beginning of each story, a white cross was displayed; after two seconds of silence, the story’s title was presented, and then the story began. The cross was always presented in the middle of the screen, and its color was randomly generated and changed every 1 to 20 seconds to keep the children’s gazes attracted throughout the story. At the end of each story, children were asked to answer an ad-hoc questionnaire (see Supplementary Materials 1.2.3). Each participant was presented with four stories, randomly drawn among the pool of ten (eleven HC participants and ten CI listened to only three stories). During the entire experimental session, their electrophysiological (EEG) activity was recorded.

### EEG recording and preprocessing

EEG data were collected, using a Brain Products system (ActiCHampPlus) with elastic caps (Easy cap Standard 32Ch actiCAP snap) for children having 32 active channels (500 Hz sampling rate). Note that for CI participants, electrodes placed very close to the magnet of the cochlear implants were disconnected (mean number of disconnected electrodes=3.50, SD=1.44; range 1–7). Continuous EEG data acquired during each story presentation were concatenated and preprocessed offline using the EEGLAB toolbox^26^, implementing a validated preprocessing pipeline^27,28^. See Supplementary Materials 1.3.1 for prototypical artifact cleaning.

### CI artifact cleaning

EEG studies involving CI users have to deal with electrical artifacts from the CI. We developed a method to clean CI electrical artifacts suited for the measurement of neural speech tracking that involves associating dynamic changes in speech features (e.g., envelope) to changes in the EEG data at specific time lags. We expected CI artifacts to occur at around 0 ms time lag^29,30^. We combined a data decomposition approach (SOBI) with an algorithm to remove components classified as containing mainly CIs artifact signals. See Supplementary Materials 1.3.2 for detailed information on the method and the discarded artifact activity.

### Filtering, removing bad channels, and epoching

After the removal of SOBI components associated with CI artifacts, (unfiltered) data cleaned from artifacts of both CI and HC groups were then low-pass filtered (cut-off=40 Hz; window type=Hanning; filter order=50), downsampled to 250 Hz, and high-pass filtered (cut-off=0.1Hz, window type= Hanning; filter order=5000). Noisy channels were identified based on the automatic bad channel detection algorithm (*clean_channels* function of *clean_data* plugin of EEGLAB; correlation threshold=0.8 and sample size=1; all the other parameters were kept as default). Noisy channels were then interpolated using spherical spline interpolation (mean interpolated electrodes per subject ± SD, in CI participants: 1.88±1.60, in HC: 2.30±1.15). Disconnected channels near the magnet of the cochlear implant were also interpolated. Following interpolation, data were re-referenced to the average reference. EEG data were then filtered according to the envelope frequency of interest: between 2 and 8 Hz (high-pass filter: cut-off=2 Hz, window type=Hanning, filter order=250, and low-pass filter: cut-off=8 Hz, window type=Hanning, filter order=126) as previously performed^31,32^. Finally, preprocessed EEG data of each story listened to was epoched (2.5 minutes, see Supplementary Materials 1.3.3 for more details), downsampled to 100 Hz, and segmented into 50-second trials, resulting in a total of 12 trials per subject (or nine for the children in which we collected three instead of four stories). Trials were created to perform a cross-validation procedure in the analysis. Data were z-scored to optimize the cross-validation procedure while estimating the regularization parameter^33^.

### Extraction of the speech envelope

For each story, the acoustic envelope was extracted, taking the absolute value of the Hilbert transform of the original piece of the story and applying a low-pass filter with an 8 Hz cut-off (3^rd^-order Butterworth filter, *filtfilt* MATLAB function).

For each subject, the speech envelope of each story was then concatenated in the same order in which they were presented to each participant and segmented into corresponding 50-second trials, resulting in twelve trials per subject (or nine trials for subjects who have listened to only three stories). The envelopes were downsampled to 100 Hz to match the EEG data^31,32^ and normalized by dividing each amplitude value by the maximum one to optimize the estimation of the regularization parameter^33^.

### Estimation of TRF

#### The forward model

To investigate how the speech envelope was tracked by the children’s brain, we used a linear forward model known as temporal response function (TRF, incorporated in the mTRF toolbox^33^). TRF can be seen as a filter that describes the mapping between ongoing stimulus features (here, envelope) and the ongoing neural response (see Supplementary Materials 1.4 for details).

We fitted separate TRF models at the single subject level to predict response in each of the 32 EEG channels from the acoustic feature (i.e., the envelope) using time lags from -100 to 600 ms in steps of 10 ms. The TRF at -100 ms time lag represents how the amplitude change of the speech envelope affects the EEG response 100 ms earlier, while the TRF at 600 ms time lag represents how the amplitude change of the speech envelope affects the EEG response 600 ms later. A leave-one-out cross-validation procedure was used to train the model. All trials except one were used to train the model to predict the neural response from the speech envelope, and the left-out trial was used to test the model. This procedure was performed for each trial; the prediction model for every trial was computed and then averaged together to obtain the TRF model for each channel.

Importantly, the regularization parameter (λ) was estimated to avoid overfitting in the regression model. The identified λ value for the envelope model was 10^4^ (see Supplementary Materials 1.4.1 for more details). These values emerged for most participants, and to generalize results, we kept λ constant across all channels and participants.

#### Estimation of the null effect (null-TRF)

To verify that neural tracking was greater than a null effect in all groups, we computed a null-TRF model for each participant^34^. We permuted the 50-second pairs of trials to obtain mismatched envelope and EEG response pairs, and then, the TRFs were fitted on these randomly mismatched trials of speech envelopes-EEG responses (*mTRFpermute* function with 100 iterations^33,35^). Then, all these null-TRF models computed across the iterations were averaged to obtain a null-TRF model that served as a control. This procedure was done separately for each participant and each channel.

#### The artifact TRF

To measure the cochlear implant electrical artifacts, we recorded the EEG activity during stimuli presentation on a phantom head^36^ with cochlear implants inserted below a conductive gel (see Supplementary Materials 1.4.2 for detailed information). TRFs were then computed on the electrical signals produced by the CIs in the absence of neural activity.

### Statistical analysis

For all analyses, the threshold level for statistical significance was set at 95% (alpha=0.05, two tails). We referred to p_FDR_ when we performed FDR correction for multiple comparisons and to p_clust_ when we performed a cluster-based permutation test.

#### Behavioral

To assess any difference in children’s comprehension, we computed the accuracy percentage (correct answers) for each participant, and we ran a univariate ANOVA with group (HC, CD, AD, HC-v) as a between-participant factor.

#### Encoding model (TRF)

First of all, we assessed the existence of neural speech tracking within HC and CI groups in a frontocentral cluster of electrodes (Cz, Fz, FC1, and FC2) typically capturing auditory responses at the scalp level^5,37,38^ and which was distant from cochlear implants. In each group, we performed comparisons between the frontocentral TRF and the frontocentral null-TRF by running paired t-tests within time lags [0–600 ms] (q=0.05, FRD correction^39^), and cluster-based permutation tests^40^ to confirm that results are stable when testing across all electrodes. The same analyses were also separately performed for the CD and AD subgroups and the HC-v group. Then we assessed whether neural speech tracking would follow a developmental trajectory in HC and CI groups. We performed a linear regression model with all the z-scored marginal moments (mean, variance, kurtosis, and skewness) of the normalized Global Field Power of the TRFs (GFP-TRF) as the independent variables and the children’s age as the dependent variable.

To investigate group differences in the spatiotemporal profiles of neural tracking, we performed cluster-based permutation tests between HC and CI irrespective of deafness onset, between CD and AD to assess the impact of the lack of auditory experience in the first year of life, and between HC-v vs. CD, and HC-v vs. AD to control for the impact of degraded speech provided by the CIs.

Finally, we investigated the relationship between neural tracking and CI’s clinical profile performing linear regression models. See the Supplementary Materials 1.5 for analysis details.

## Data availability

Since participants are minors, we cannot share original raw data, but individual TRF and statistical code analysis are shared in this Mendeley repository doi: 10.17632/nzg5g2gzrd.2.

## Results

### Neural speech tracking in hearing and cochlear-implanted children

First, we assessed whether neural speech tracking could be measured in HC and CI children irrespective of deafness onset. We estimated the temporal response function (TRF) within a frontocentral cluster of sensors suitable for measuring auditory response functions in children and adults^5,41^ and in CI users^37^. Contrasting the TRF model with the null-TRF in both HC and CI children, we observed significant neural speech tracking (see Figure 2A). Specifically, in HC children, auditory responses were characterized by a prominent positivity between 0 and 110 ms time lags (all p_FDR_<0.05; peak TRF=0.065, SE=0.008; d=1.24, 95^th^ confidence interval (CI95)=0.78–1.64, see Supplementary Materials 2.1 for details on how Cohen’s d was computed), and second negativity between 200 and 320 ms time lags (all p_FDR_<0.05; peak TRF=-0.051, SE=0.009; d=-0.86, CI95 =-1.21–-0.39). In the CI group, the first positive response emerged at time lags between 20 and 270 ms (all p_FDR_<0.05; peak TRF=0.089, SE=0.010; d=1.92, CI95=1.43–2.49), and then the negativity occurred after 390 ms time lag (all p_FDR_<0.05; peak TRF=-0.054, SE=0.007; d=-0.93, CI95=-1.44–-0.48). Results highlighted that in both groups of children (i.e., HC and CI), the neural tracking could be robustly measured with twelve minutes of natural speech; the activity being characterized by two main phases of short and long timescales of brain-speech tracking occurring within 600 ms (see Supplementary Materials 2.2 for confirming results across all electrodes Figure S1, and Figure S2 for visualization of the whole spatiotemporal dynamics).

**Figure 2.**
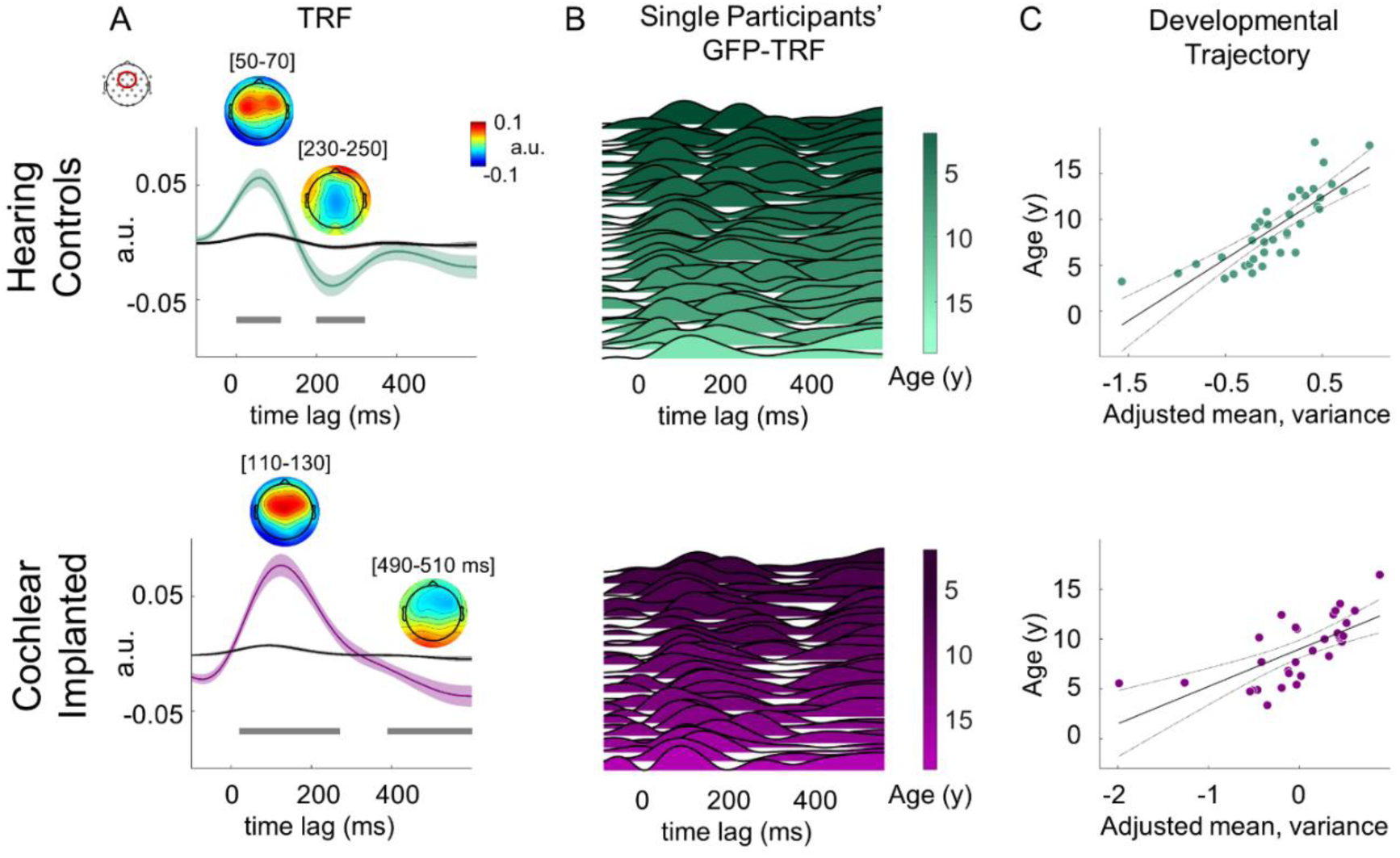
Neural speech tracking in HC and CI children. (A) Grand average speech tracking TRFs (olive color HC, magenta color CI) with topographies to represent peak distribution over the scalp and grand average null-TRFs (grey color) at frontocentral electrodes (Cz, Fz, FC1, and FC2) between -100 and 600 ms time lags. Shaded areas represent SE of the mean. Grey horizontal bars indicate time lags (between 0 and 600 ms) at which speech-tracking TRFs significantly differed from the null-TRF (t-test, FDR corrected pFDR<0.05). (B) Single participants’ Global Field Power of their speech tracking TRF (GFP-TRF, normalization was performed for visualization purposes) sorted by age; HC group in the upper panel, and CI group in the lower panel. (C) Partial regression plots of the linear regression models show a developmental trajectory in both HC (upper panel) and CI (lower panel) groups (variance and mean of the GFP-TRF were associated with children’s age).

Once the existence of the speech-tracking TRF was verified in both groups, we investigated whether it was possible to measure a developmental trajectory of neural tracking in both HC and CI children. This would strongly advocate for employing neural speech tracking as a reliable measure even in children who perceive hearing only through cochlear implants. We reasoned that, with age, neural tracking would become more efficient. Therefore, the TRF, representing the synchronization between the neural signals and the continuous speech, would become less spread over time as typically observed in developmental ERP and TRF studies^5,42^ and as suggested by energy landscape analysis methods applied to EEG signals^43,44^. In turn, we hypothesized that with increasing age, the TRF amplitude would be condensed in fewer time lags, and more time lags would have no substantial neural tracking. In other words, we expected the signal’s sparsity to increase with age as the TRF would have a higher variance of values over time. To this aim, first, we computed at the single participant level the Global Field Power (GFP^45^) of the TRFs (i.e., GFP-TRF) for a better estimate of the dynamic of the signal and to avoid a space-dependent index^46^. For each GFP-TRF between -100 and 600 ms, we estimated the marginal moments (i.e., variance, mean, kurtosis, and skew) to characterize the data distributions, and we tested with a linear model whether the estimated marginal moments of the GFP-TRF were associated with children’s age. In the HC group a clear developmental pattern emerged, highlighting an association between neural tracking signal and age, with an increase of variance (signal sparsity) and a decrease of mean (adjusted R^2^=0.62, F_(4,32)_=15.7, p<0.001; variance: β=3.27, SE=0.95, p=0.002; mean: β=-5.89, SE=0.91, p<0.001, see Figure 2C upper panel). A similar pattern was observed in CI children (adjusted R^2^=0.42, F_(4,27)_=6.6, p<0.001; variance: β=2.08, SE=0.68, p=0.005; mean: β=-3.11, SE=0.65, p<0.001, see Figure 2C lower panel).

### How temporary auditory deprivation affects neural speech tracking

Once the existence and development of the auditory response function were assessed in both HC and CI children, we compared the TRFs across groups. First, spatiotemporal profiles of TRFs were compared between the CI and the HC groups using a cluster-based permutation test performed at the whole brain level (across all sensors) and comprising the TRFs at every time lag between 0 and 600 ms. Results revealed a significant difference between CI and HC groups (p_clust_<0.05), with the largest significant effect at time lags between 110 and 290 ms in a large frontocentral cluster of sensors (d=-1.69, CI95=-2.14–-1.25). This difference revealed that neural tracking in children with CIs was delayed, already at a short timescale of brain activity, at the first TRF peak, and the subsequent activity was hampered (see Figure 3A).

**Figure 3.**
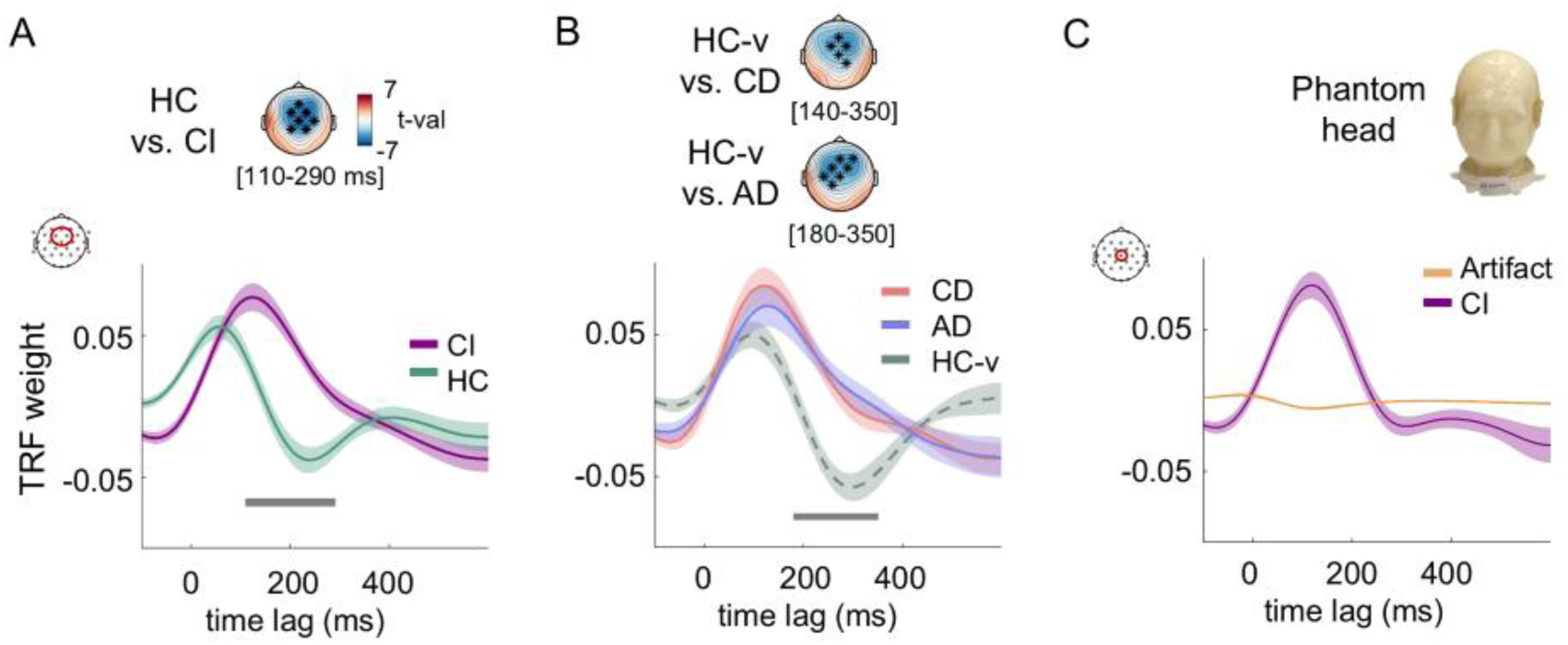
The impact of auditory deprivation on neural speech tracking. (A) The speech tracking TRF at the frontocentral sensors between -100 and 600 ms time lags for HC (olive) and CI (magenta) groups. The topography shows the statistical difference between TRFs in HC and CI children between 110 and 290 ms; significant sensors at pclust<0.05 are highlighted with black asterisks. (B) Speech tracking TRF at the frontocentral sensors between -100 and 600 ms time lags for CD (red), AD (blue), and HC-v (dashed greenish) groups. For all TRFs, the continuous line represents the group mean and the shaded area the SE. The data of CD and AD overlapped (no difference in the cluster-based test), suggesting that the auditory experience in the first year of life does not affect neural speech tracking in CI children. Both CD and AD are significantly different from HC-v groups (significant sensors at pclust<0.05 are highlighted with black asterisks), unveiling that the anomalies in CI neural tracking cannot be explained merely by the vocoded speech. (C) Artifact and CI’s TRF are shown for the electrode of the frontocentral cluster where the artifact activity is stronger (Cz). Results suggest negligible impact of the electrical activity of the implant on the TRF measured in CI children.

Next, we investigated the specific role of auditory experience in the first year of life. The two groups of CI children, with congenital deafness (CD) and acquired deafness (AD), were contrasted to investigate the role of perinatal auditory input in the development of neural speech tracking (note in both CD and AD groups the TRF exceeded the null-TRF at frontocentral sensors; see Supplementary Materials 2.3 and Figure S3A and B). The data revealed a clear overlap between the TRFs of the two CI groups (see Figure 3B). No difference emerged between CD and AD (cluster-based permutation test performed across all sensors between 0 and 600 ms, no clusters were found at p_clust_<0.05). This suggested that the alterations of neural speech tracking observed in CI compared to HC occurred irrespective of whether children experienced or not the lack of hearing input within the first year of life (see Supplementary Materials 2.4 for a contrast showing that the TRF of both CD and AD groups differed from the HC group).

We further assessed whether the difference between HC and CI’s neural tracking could be driven by the degraded stimulation provided by the cochlear implants compared to the natural sounds perceived by HC children. Similarly to the comparison with HC, significant differences emerged between hearing children who listen to vocoded-speech (HC-v) and both CD and AD groups (p_clust_<0.05), indicating that alterations in CI’s neural tracking cannot be explained by the mere degradation of the speech signal (Figure 3B).

Finally, since cochlear implants work by producing electric impulses, we assessed whether functioning cochlear implants could profoundly affect the recorded and modelled EEG signal. We therefore measured the artifact TRF obtained from the electrical activity of cochlear implants during stimuli presentation to a phantom head in the absence of brain activity. Artifact TRF had negligible magnitude and, compared to CI’s TRF, had a completely different profile, with the first peak at 0 ms as expected (Figure 3C). This observation ruled out the possibility that the implant activity merely caused alterations emerging in the neural speech tracking of the CI group.

To sum up, results demonstrated that the difference between CI and HC cannot be explained by the absence of auditory experience during the first year of life, by degradation of the speech signal, or by implant activity. Given that neural tracking of CD and AD children did not differ, we further characterized the alterations of the neural speech tracking of CI children as a single group and the association of these alterations with clinical profiles aiming to unravel possible biomarkers of their continuous speech processing.

The first anomaly of CI’s neural tracking is the latency. Data revealed that the first peak (i.e., GFP-TRF P1) in the CI group was substantially delayed compared to the HC group (t_(67)_=-2.97; p=0.004, HC mean=86.8 ms, SE=8.7; CI mean=119.1 ms, SE=6.0; d=-0.71, CI95=-1.24–-0.12, see Figure 4A), suggesting less efficient neural tracking over short time intervals from sound onsets. Given that previous neurophysiological studies^23^ have highlighted the impact of implantation age on auditory response latency to simple sounds, we assessed whether the age at implantation affects TRF latency. A linear regression indicated that the age at which auditory input was restored with cochlear implantation accounted for the delayed neural tracking in CI users (latency of the neural tracking first peak, GFP-TRF P1). Data indicated that the later the implantation occurred, the more delayed the neural tracking (R^2^=0.115, F_(1,30)_=5.04, p=0.032, β=0.38, CI95=0.04–0.72, see Figure 4B). Note that other clinical variables such as chronological age, age at deafness onset, age at which hearing aids were provided before implantation, and experience with the implant, did not explain the delay (p>0.22). Given the high variability of the neural tracking first peak (GFP-TRF P1) for the earliest implanted children (see Figure 4B), we further investigated the association between implantation age and TRF latency by fitting multiple linear functions to identify discontinuities (Friedman, 1991 and see Supplementary Materials 2.5). We found a significant regression model with two basis functions and a discontinuity knot at 21 months (R^2^=0.173; cross-validated R^2^_GCV_: 0.06, p=0.036, hinge function max (0, x1 -21) β=0.45, CI95=0.44–1.05), revealing that the positive relationship between implantation age and the latency of the neural tracking emerged from 21 months onward. Note that the other clinical variables cannot explain the GFP-TRF P1 variability before 21 months of age (see Supplementary Materials Figure S4).

**Figure 4.**
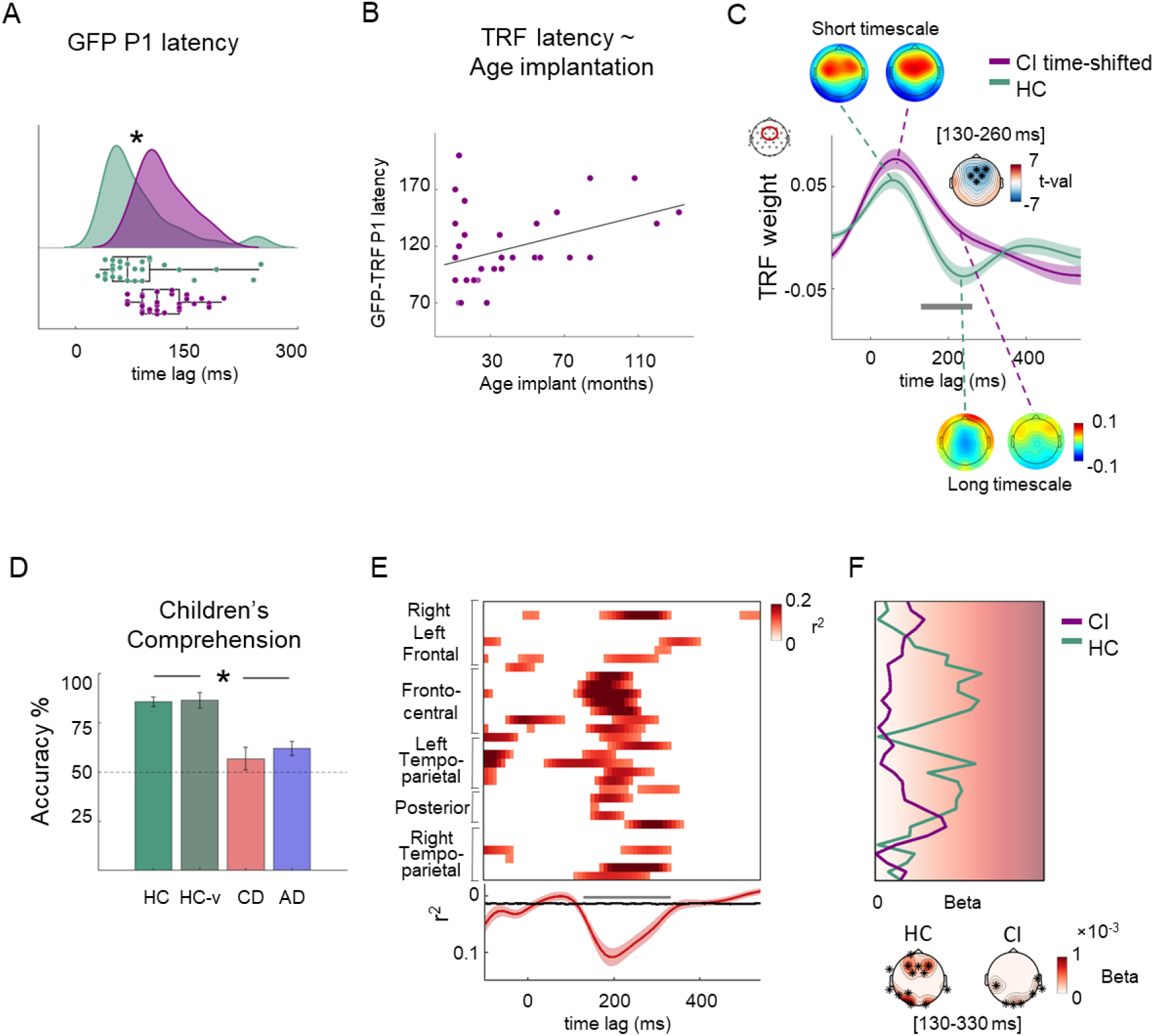
Biomarkers of altered neural speech tracking in children with CIs. (A) The first peak latency (GFP-TRF P1) was delayed in CI compared to HC (t(67)=-2.99; p=0.004). (B) Linear regression predicting the neural tracking first peak (GFP-TRF P1) from the age at which CI children received the first implant (auditory restoration). (C) The plot shows the TRF for HC and CI at frontocentral sensors. The first peak of the CI’s TRF was temporally realigned to the first peak of the HC group to account for the CI’s neural tracking delay. The topographies show the similarity at a short timescale (P1TRF), while a clear difference at a longer timescale (N2TRF). After accounting for the delay, significant differences between the two groups emerged selectively in the time window [130–260 ms]; significant sensors are highlighted with black asterisks. (D) Comprehension scores in HC, HC-v, CD, and AD groups. Children with CIs (both CD and AD) had lower scores compared to both HC and HC-v groups. (E) The plot shows the r^2^ values with p<0.05 for the regression model in which the point-by-point TRF values were predicted by behavioral accuracy, group, the interaction between them, and age. Below, the r^2^ averaged across sensors is plotted as a function of time, and the shadow represents the SE. The black line represents the boundary of the null effect, that is, the 95th percentile of the r^2^ null distribution (note that the y-axis is reversed; higher averaged r^2^ values emerged below the black line). A significant difference emerged between 130 and 330 ms (pFDR<0.05). (F) The plot shows the absolute values of beta accuracy averaged within the significant time window [130–330 ms] across all sensors (neighboring channels are smoothed for visualization purposes), separately for HC and CI groups. Below are shown the topographies with black asterisks highlighting significant sensors for each group. Data revealed the relationship between TRF magnitude between 130 and 330 ms and children’s comprehension scores in both groups. CI group had markedly reduced neural speech tracking at this latency, and a reduced association between comprehension scores and TRF magnitude.

A second anomaly of the CI’s neural speech tracking emerged at a longer timescale: a difference between the TRF dynamic of CI and HC children was evident even when accounting for the neural tracking delay (see Figure 3A and 4C). To test this hypothesis, we shifted the CI’s data, aligning the first TRF peak (P1, at the frontocentral cluster) of the CI group to the homolog peak of the HC group (see Supplementary Materials 2.6 for details). The cluster-based permutation test (across all sensors and time lags) between HC’s TRFs and CI’s temporally shifted TRFs revealed reduced neural tracking in CI individuals compared to HC at time lags between 130 and 260 ms (p_clust_<0.05; d=-1.11, CI95=-1.50–-0.69). At this long timescale of brain activity, the second major phase of the auditory temporal response function (N2_TRF_) emerged clearly only in the HC. Conversely, the first TRF phase (P1_TRF_) did not differ between HC and CI, suggesting unaltered TRF magnitude at a short timescale (Figure 4C).

### Neural tracking and speech comprehension

To assess speech comprehension, we investigated the outcome of the behavioral questionnaire, which comprised questions concerning story details. Results revealed impaired scores in CI children compared to HC (one-way ANOVA: F_(3,81)_=16.96, p<0.001). Both CD and AD groups showed significantly lower accuracy with respect to HC and HC-v children (HC: mean 85.70 % accuracy ± SE 2.50; HC-v: mean 86.46 % accuracy ± SE 3.87; CD: mean 56.77 % accuracy ± SE 5.72; AD: mean 61.98 % accuracy ± SE 3.61). Post-hoc Bonferroni corrected t-tests confirmed significant differences between both HC and CI groups (HC vs. CD p_Bonf_<0.001, d=1.60, CI95=0.93–2.31; HC vs. AD p_Bonf_<0.001, d=1.56, CI95=0.83–2.34; HC-v vs. CD p_Bonf_<0.001, d=1.48, CI95=0.81–2.13; HC-v vs. AD p_Bonf_=0.001, d=1.60, CI95=0.56–2.45, see Figure 4D), while no difference emerged between the two CI groups nor between the two HC groups (p_Bonf_=1.00).

Following previous evidence linking neural speech tracking and comprehension, we tested whether the degree of neural tracking (the TRF magnitude) would be associated with children’s comprehension scores^47^ by performing a series of linear regression models (at all sensors) to predict the point-by-point TRF values from response accuracy, groups, and their interaction, accounting for the impact of participants’ age (see Supplementary Materials 2.7). The model was significant between 130 and 330 ms (averaging across all sensors, p_FDR_<0.05, mean r^2^=0.073, SE=0.009, see Figure 4E and S5A for detailed results). When we computed the regression model separately for each group, beta accuracy (averaged between 130 and 330 ms) revealed a significant impact of children’s comprehension on TRF magnitude at both frontal and posterior sensors in HC children. Instead, in the CI children, this relationship was less strong and emerged only at posterior sensors (see Figure 4F and S5B and C for more details). Overall, these findings revealed that the magnitude of the TRF second phase (N2_TRF_ between 130 and 330 ms), i.e., long timescale neural tracking, was associated with children’s comprehension scores. Precisely at this latency (note, once accounting for their neural tracking delay by time-aligning the CI’s and HC’s TRFs) the CI group had markedly reduced neural speech tracking. This altered dynamic accounted for their comprehension deficit.

## Discussion

Despite the period of auditory deprivation and the fact that cochlear implants provide only partial input to the brain^12,13^, our data clearly revealed the possibility of measuring neural speech tracking in CI children and the informativeness of the associated indices.

First, our data revealed a developmental trajectory in the CI group similar to the HC group, supporting the sensitivity of neural tracking to capture functional changes in both typical and atypical development (see ^6,48^). Most importantly, the TRF remarkably overlapped in individuals with congenital deafness (CD) and acquired deafness (AD). Moreover, at a short timescale, the neural speech tracking had similar magnitude between HC and children with CIs. This evidence favours the model in which the brain is endowed with biological constraints for the tracking of speech that are resilient to auditory deprivation in the first year of life, expanding previous observations that certain basic computations of language tracking are already available at birth^49^.

When we contrasted the neural tracking between CI and HC groups, we observed two main differences: the CI’s neural response was delayed with respect to HC at a short timescale (P1_TRF_) and dampened at a longer timescale (N2_TRF_). Importantly, these anomalies in CI’s TRF dynamics were related to biological differences in brain processing: they were not merely caused by degraded speech stimulations, nor by the impact of the CI electrical response artifact. Indeed, neural tracking was similarly affected in CI compared to hearing children, irrespective of whether the latter were exposed to natural speech or vocoded-speech. Moreover, the artifact TRF computed from cochlear implants in the absence of brain activity could not account for the main results.

### Delayed neural tracking in children with CIs

While at a short timescale, the neural speech tracking had similar magnitude, it was markedly delayed in the CI children compared to HC individuals (of about 60 ms). This is coherent with previous observations employing simple speech units, such as syllables^23,25^. Interestingly, in adults, the severity of hearing impairment is positively associated with neural tracking delay^50^. Coherently, difficult acoustic listening conditions, such as when speech is presented with background noise, are known to delay auditory responses to speech stimuli measured with ERPs or neural tracking in both children and adults^51–54^. Noteworthy, hearing children who listen to vocoded-speech showed a slightly delayed P1_TRF_ (100 ms, see Figure S3C) with respect to those who listened to normal speech (60 ms). Taken together, these findings might suggest that delayed neural tracking observed in children using CI could reflect a reduced efficiency of continuous speech processing.

Crucially, the TRF delay measured here was associated with the age of cochlear implantation. Coherently with seminal electrophysiological studies employing short-lived speech sounds^21–24^, our results highlighted the crucial role of age at which auditory restoration takes place for the development of speech tracking. The later the child receives the implant, the more delayed is the first TRF peak (P1_TRF_), which represents auditory processing of language at a short timescale (within the first 150 ms of brain-speech tracking). Longitudinal studies on cochlear-implanted children established a sensitive period for the development of basic auditory responses to syllables within the first 3.5–4.0 years of age^55,56^, strongly advocating for early implantation. In the case of children implanted at an early age (< 3.5 years), the P1 latency of the ERPs consistently fell within the 95% confidence interval of typical development. Conversely, children who underwent implantation after the age of 7 years never reached the typical latency range of early auditory responses^23,57^. Here, we found further support for these observations, showing experience-dependent effects associated with the timing of implantation, revealing that they also seem to emerge for early, sensory-based components of the neural tracking of continuous speech. Importantly, our data shows that the benefit of early implantation was evident only from 21 months of age.

### Neural tracking dynamics uncover higher-order deficits of speech processing in children with CIs

The neural tracking magnitude occurring at a long timescale of brain activity (N2_TRF_, between 130 and 330 ms) was associated with higher comprehension scores. This result is consistent with recent evidence suggesting that the magnitude of neural tracking with a similar timescale is associated with comprehension of continuous speech^58^. Coherently, studies using noise-vocoded speech demonstrated that neural tracking occurring at about 200 ms was strongly reduced when the speech was degraded and comprehension impaired^59^. Even accounting for the delay, by time-aligning CI’s and hearing control’s neural data, CI individuals had hampered tracking at this long timescale (see Figure 4B). Coherently, the behavioral performance of both CD and AD groups was markedly impaired, and the association between neural tracking magnitude and behavioral performance was less strong in CIs compared to HC. These findings provide the first evidence for the identification of a possible biomarker associated with natural speech comprehension in CI individuals. TRF magnitude at this latency range (N2_TRF_) could be employed to verify children’s speech understanding especially when behavioral measures are difficult to acquire, as in the case of infants, and to estimate the development of speech processing after implantation.

Finally, it is important to stress that our findings are clearly biologically driven: (i) a developmental trajectory similar to hearing peers emerged in CI groups, (ii) neural tracking anomalies of CI children could not be ascribed to the degraded stimulation cochlear implant convey nor to their electrical artifacts (iii) the latency of CI’s neural tracking was affected by the age at which children received the implant, and finally, (IV) the dampened neural tracking occurring at long timescale was linked with their impaired speech comprehension. Taken together these results revealed that altered CI’s TRF dynamics are the result of their atypical auditory experience.

### Limitations

Despite results pointing toward the crucial role of early auditory restoration in mitigating atypical auditory development and possibly ameliorating speech processing efficiency, we cannot provide conclusive interpretations to what causes the high variability of neural tracking latency (GFP-TRF P1) observed across children implanted before 21 months of age (latency from 70 to 200 ms). The aetiology of children implanted before 21 months of age comprised different profiles (e.g., GJB2 connexin 26, congenital CMV, Waardenburg syndrome, and perinatal complications associated with prematurity). Most of them had bilateral profound congenital deafness onset (ten out of eleven, see Supplementary Materials Figure S4B). Noteworthy, in the case of congenital deafness onset, the clinical practice does not assess whether auditory input was available in the intrauterine life since auditory screening is performed only after birth^60^. Intriguingly, recent evidence suggests that fetal linguistic experience shapes neural synchronization with language measured at birth^61^.

In conclusion, the data clearly highlighted that speech tracking leverages a robust biological predisposition, resilient to a period of auditory deprivation from birth, strongly supporting treatment opportunities available for children with congenital deafness. However, we substantiated in CI children a specific vulnerability of higher hierarchical levels of neural speech tracking, which are associated with speech comprehension. Thanks to its simple nature, neural speech tracking is a promising method to assess auditory and speech functions in developing cochlear-implanted individuals and could help explain the variability in outcomes that typically characterize this population.

## Abbreviations

CI: Cochlear Implant
CD: Congenital Deafness
AD: Acquired Deafness
HC: Hearing Control
TRF: Temporal Response Function.

## Acknowledgments

We want to acknowledge all the children and their families participating in this study.

## Author Contributions

Conceptualization: DB, AF

Methodology: AF, DB, GH, EB, MB, MF

Investigation: AF, MF, EN, AM, DB

Visualization: DB, AF, GH

Funding Acquisition: DB, ER, FP

Project Administration: DB

Supervision: DB

Writing – Original Draft: DB, AF

Writing – Review & Editing: DB, AF, GH, FP, EN, EO, ER, MF, MB, BB, AM, EB

## Supplementary Materials

Supplementary material is available at Brain online.

## Supplementary Materials

### 1 Materials and Methods

#### 1.1 Participants

##### 1.1.1 Sample size estimation

We estimated the minimum sample size needed to measure a clear P1 in the auditory response function. We expected the P1 to emerge between 0 and 150 ms lags ^1^ and in the frontocentral sensors (Fz, Cz, FC1, FC2), and it should be higher than in the null-TRF. Using the data of 10 pilot HC subjects, we computed the mean and SD acoustic TRF (mean = 0.044, SD = 0.036) and null-TRF (mean = 0.005, SD = 0.007). Through simulations suited for cluster-based permutation tests (500 randomizations^2^), we estimated a minimum sample size of 16 subjects to reach a power of 0.95 (lower threshold).

##### 1.1.2 Participants exclusion

Five CI participants were excluded due to different reasons: (i) a two-year-old child was discarded because they could not comply with the experimental session; (ii) two children were excluded because their IQ was below the age standard; iii) one child was reimplanted after many years from the first implantation following an ear infection; iv) one child was discarded due to the bad quality of the EEG signal (15 electrodes were detected as bad channels). The remaining thirty-nine CI participants were classified according to their deafness onset: children with profound bilateral congenital deafness (CD) and children who acquired profound bilateral deafness during the development (AD). Since it was not possible to ensure whether profound bilateral deafness was congenital or acquired in seven children, their data were not analyzed.

##### 1.1.3 Criteria for congenital vs acquired deafness classification

Congenital deafness (CD) was ensured by the following criteria: (a) having failed to pass the neonatal screening for otoacoustic emissions (typically < 1 week after birth); (b) receiving a diagnosis of profound bilateral deafness (hearing thresholds ≥ 90 dB in both ears) following the objective evaluation of auditory brain-stem responses (ABR) within two months of age (median age: 36 days IQR = 21.5; range 21 – 60 days). In contrast, AD children had at least some auditory experiences in early development (i.e., a minimum of 12 months). To ensure the presence of such auditory experience, we combined the following clinical information: (a) whether they passed otoacoustic emissions neonatal screening at least with one ear; (b) an ABR indicating normal hearing before the diagnosis of deafness or a diagnosis of deafness that was not profound bilaterally (e.g., moderate deafness at least in one ear) made by ABR or behavioral test (hearing thresholds < 90 dB in at least one ear); (c) family report indicating residual hearing for the first period of life. All patients with acquired deafness (AD) received a diagnosis of profound bilateral deafness before cochlear implantation (hearing thresholds ≥ 90 dB in both ears for children under 2 years old and > 75 dB for children older than 2 years old ^3^), age range: 12 – 107 months. If the exact date of this test was unavailable, we estimated the onset of profound bilateral deafness one month before the date of the first cochlear implantation; N = 6).

##### 1.1.4 Ethic committees

Comitato Etico Regionale per la sperimentazione clinica della Regione Toscana: Numero registro 34/2020, and Comitato etico congiunto per la ricerca espressione di parere delibera n. 17/2020.

#### 1.2 Stimuli and Procedure

##### 1.2.1 Stimuli registration, preprocessing and presentation

Stories were read by a person whose diction had been formally trained and were recorded in a sound attenuated chamber (BOXY, B-Beng s.r.l., Italy) with an iPhone 7 (camera with 12MP, video resolution in HD, 720p with 30fps, at a sampling frequency of 48000 kHz) and an external condenser microphone (YC-LM10 II, Yichuang). All audio recordings were imported in iMovie (version 10.3.1), the noise reduction at 100% was applied, and each file was cut to have 2 seconds of silence before the story’s title and a few seconds of silence at the end of the story. Then, the audio was imported into Audacity® (version 2.4.2, https://www.audacityteam.org/ using ffmpeg and lame functions to isolate the audio from the video). The audio files imported in Audacity were preprocessed with the following steps: they were converted from stereo to mono, amplified (default value in Audacity and avoid clipping were selected), down-sampled to 44100 Hz, and set to a 32-bit sample. Finally, we imported all audio files in Matlab to perform RMS equalization to achieve an equal loudness for all the stimuli (RMS value = 0.03). Stimuli were presented to participants using Psychopy® software (PsychoPy3, v2020.1.3).

##### 1.2.2 Vocoded version of the speech stimuli

To simulate acoustically what CI users experience through their devices, we created a vocoded version of our stories with the vocoder function (https://github.com/egaudrain/vocoder), applying the frequency spacing as in Cochlear devices and implementing the ACE processing strategy.

##### 1.2.3 Comprehension questionnaire

Each questionnaire at the end of the story comprised two comprehension questions, 2-alternative-forced-choice; most of them were yes-no questions (e.g., “Did Lorenza like to travel?”) and few alternative answer questions for the young children (e.g., “What did the kitten fairy give to Lorenza?”, possible answer in the picture “ball of wool” or “doll”). For younger children (age range between 3 – 6 years), questions were performed verbally by the narrator’s voice, and the alternative answers were supported by drawings representing the content. For older children (> 7 years old), questions were presented via text on the screen and read by the experimenter.

#### 1.3 EEG preprocessing

##### 1.3.1 Prototypical artifact cleaning

Continuous EEG recordings were low-pass filtered (cut-off=40 Hz; window type=Hanning; filter order=50), downsampled to 250 Hz, and high-pass filtered (cut-off=1 Hz; window type=Hanning; filter order=500). The filtered downsampled data were segmented into consecutive 1-second epochs. Noisy segments were removed using joint probability (threshold across all channels=3 SD ^4^). To remove prototypical artifacts (e.g., blink and eye movements), data were submitted to Independent Component Analysis (ICA, based on the extended Infomax ^5–7^. The computed ICA weights were applied to the continuous *raw* (unfiltered) data ^8,9^. Components associated with blinks and eye movement artifacts were identified using CORRMAP, a semiautomatic procedure in which a prototypical topography for each type of artifact (i.e., eye movement and blink) is selected. All the components that correlate more than 80% with the template were removed ^10^. For CI participants, the mean number of removed components was 2.09±0.39 SD, and for HC, 2.00±0.00 SD.

##### 1.3.2 CI artifact cleaning method

To search for CI activity that could generate artifacts in the EEG data, we performed the following steps. We decomposed the EEG recordings into components (using the SOBI algorithm) with the purpose of separating physiological and noise sources. Then, we applied the TRF approach to each component to obtain a set of component-TRFs. Finally, to identify artifacts with about zero lag, and by using a minimal set of parameters extracted from TRFs in HC, we identified and discarded the CI artifactual SOBI components and reconstructed back EEG recordings. These steps are explained below in detail.

Data cleaned by their stereotypical artifacts (blinks and eye movements) were preprocessed using the second-order blind identification (SOBI) algorithm to identify independent components based on second-order statistics, making it suitable to separate temporally correlated signals and maximizing activity related to CI artifacts ^11^. The same procedure explained above for ICA (filtering and rejection of noisy segments) was applied prior to SOBI estimation. Specifically, continuous cleaned EEG recordings were low-pass filtered (cut-off = 40 Hz, window type = Hanning, filter order = 50), downsampled to 250 Hz to reduce the computational time, and high-pass filtered (cut-off = 1 Hz, window type = Hanning, filter order = 500). The filtered downsampled data were segmented into consecutive 1-second epochs. Noisy segments were removed using joint probability (threshold across all channels = 3 SD; ^4^. Then, SOBI was computed, and the SOBI weights were applied to the original (unfiltered) data cleaned by their stereotypical artifacts. Data were downsampled to 250 Hz, filtered between 2 – 8 Hz, epoched from 6 seconds to 2.5 minutes for each story, downsampled again to 100 Hz in order to match the sampling rate of the acoustic features for TRF estimation, and segmented into 50-second trials (see the following paragraph “Filtering, Removing Bad Channels and Epoching” for a more detailed explanation of these steps). The TRF model was applied to each SOBI component across -100 – 600 ms time lags (the same parameters were used for the envelope TRF encoding model; see below). Thus, we obtained a TRF of each SOBI component for each subject.

Subsequently, we implemented an algorithm to remove the components classified as containing mainly CIs artifact signals. We started defining the criteria to reject components using the data of the HC participants (who have no CI) to identify parameter values at a 5% false positive rate. First, we normalized SOBI components by extracting the absolute value of each timepoint and scaling it with the maximum intensity of the series. Then, we modeled normalized SOBI components by fitting in each time series a set of Gaussian responses: (i) a single Gaussian was restricted to peak between -100 and 0 ms (that should contain mainly artifactual activity); (ii) up to five Gaussians were limited to peak between 50 and 500 ms (that should include mainly neural activity). Two criteria were used to decide whether a SOBI contained artifacts: (a) the *ratio* between R^2^ of the Gaussian fitted before zero and R2 Gaussians fitted after zero; (b) the beta of the Gaussian fitted before zero. The rationale was that we expected to find most implant activity around zero, while the rest (after a physiological delay) should be considered brain activity. In HC, we mapped our R^2^ *ratio* and *beta* onto a plane to identify a decision boundary that isolates portions of the parameters space with both high *R*^2^ *ratio* and *beta* and retains a false positive rate of 5%. This decision boundary was applied to CI data using the same procedure described above. Artefactual components with high R^2^ *ratio* and *beta* were removed (number of components removed per participant mean ± SD: CI 2.47 ± 2.19, CD: 2.69 ± 2.39, and AD: 2.25 ± 2.02).

##### 1.3.3 Epoching details

The timing of each epoch was adjusted to +99 ms onset delay measured by the AV device (EGI). Then, preprocessed EEG data of each listened story were epoched starting from 6 seconds and lasting 2.5 minutes. The first 6 seconds of each story, including 2 seconds of silence, the story title, and the beginning of the story, were removed to avoid the stimulus onset response as much as possible ^12^. Accordingly, for each audio file the first 6 seconds were discarded.

#### 1.4 The temporal response function (TRF) model

The forward model approach allows the prediction of previously unseen EEG responses from the stimulus feature and has been extensively used to model the neural tracking of continuous speech envelope. Mathematically, the encoding model is described by the following function:

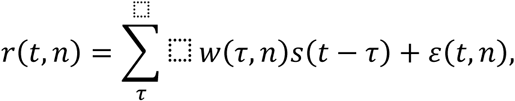

Where *t* = 0, 1, … T is time, *r(t,n)* is the neural EEG response from an individual channel *n* at time *t*, *s* is the stimulus feature(s) at each moment (*t – τ*), τ is the range of time lags between *s* and *r*, *w(τ,n)* are the calculated regression weights over time lags (TRF), and *ε(t,n)* is a residual response at each channel *n* at time *t* not explained by the TRF model ^13^. Specifically, the TRF at each time lag (τ) represents how the unit change in the amplitude of the speech envelope would affect the EEG response τ ms later ^14^.

Importantly, since the TRF model is conducted separately for each channel, their interpolation during preprocessing (in case of bad channels or vicinity with the cochlear implant) did not affect the model results.

##### 1.4.1 Regularization parameter estimation

Overfitting consists of fitting the noise in the data unrelated to the stimulus, thus preventing generalization to different datasets. Regularization is achieved by selecting the TRF models’ optimal regularization parameter (λ). A set of ridge values (λ = 10^-3^, 10^-2^, 10^-1^, 1, 10, …, 10^9^, 10^10^) is used to compute the model for time lags from -100 to 600 ms through a leave-one-out cross-validation procedure. To determine the optimal regularization parameter (λ) for each participant, we used the mean squared error (MSE) value – averaged across trials and electrodes – between the actual and the predicted EEG responses; the λ value reaching the lowest MSE value was selected.

##### 1.4.2 The artifact TRF measured on EEG phantom head

We created our EEG phantom head following the information from the Open EEG Phantom Project ^15^. We inserted two cochlear implants below the phantom “skin” made of ballistic gelatin, and then we applied the EEG on the phantom head to record electrical activity during stories presentation. We recorded three different sessions, one with both cochlear implants switched on, one only with the left, and one only with the right implant operative. During each of these sessions a total of twelve stories were presented (four for each group of ages 3-6, 7-10, and 11-15).

The EEG data were preprocessed following the same steps implemented for participants except for the removal of eye movement and blink artifacts, and then TRF was computed.

#### 1.5 Statistical analysis

##### 1.5.1 Assessing the existence of neural speech tracking (TRF) within each group

To test whether we could measure an auditory temporal response function (TRF model) within HC and CI groups, we first selected a cluster of four frontocentral channels (Cz, Fz, FC1, and FC2; note that none of these channels were disconnected in CI children) typically capturing at the scalp level auditory responses with evoked potentials (for review see ^16^) and auditory response functions (TRF) in children and adults (e.g., ^11,17^). Then, we performed comparisons between the frontocentral TRF model and the frontocentral null-TRF by running paired t-tests within time lags [0 – 600 ms] (i.e., every 10 ms) separately for HC and CI groups and also separately for CD and AD subgroups, and for HC-v group (two-tailed, q=0.05, FRD correction ^18^). We also performed cluster-based permutation tests ^19^ in the FieldTrip toolbox ^20^ between the TRFs and the null-TRFs within each group to confirm that the same results emerged testing the frontocentral cluster are stable also testing across all electrodes. A cluster was defined along electrodes × time lags dimensions. Cluster-based permutation tests were performed at the whole brain level (across all electrodes) and time lags between 0 and 600 ms, using the Monte-Carlo method with 1000 permutations. Cluster-level statistics were calculated by taking the sum of the t-values within every cluster (minimum neighboring channel = 2; cluster alpha was set to 0.05, which was used for thresholding the sample-specific t-statistics). Identified clusters were considered significant for the permutation test at p < 0.025 (the probability of falsely rejecting the null hypothesis). The alpha level 0.05 was thus divided by 2 (p = 0.025) to account for a two-sided test (positive and negative clusters).

##### 1.5.2 Investigating the developmental trajectory separately for the HC and CI groups

We assessed whether neural speech tracking would follow a developmental trajectory in HC and CI groups. We expected the TRF signal to become sparser with age. In this case, sparsity would indicate selectivity (in the time domain) of natural activity. This would result in high-variance amplitude distributions. To this aim, we estimated marginal moments of the TRF signal as they allow us to describe the sparsity of a signal (e.g., ^21,22^). We performed a linear regression model with all the z-scored marginal moments (mean, variance, kurtosis, and skewness) of the normalized Global Field Power of the TRFs (GFP-TRF) as the independent variables and the children’s age as the dependent variable. The reason for choosing the GFP instead of selecting specific channels of interest is that this approach allows for a more objective and reference-free characterization of temporal dynamics of the global electric field (see ^23^).

##### 1.5.3 Testing neural tracking differences between groups

To test differences in the spatiotemporal profile of TRFs between groups, we performed cluster-based permutation tests with the same parameters defined above. First, we performed a cluster-based permutation using independent-sample t-statistics between HC and CI groups without any a priori hypothesis on time or space. Then, we performed the same cluster-based permutation test between CD and AD to investigate the impact of the lack of auditory experience in the first year of life, and between HC-v vs. CD, and HC-v vs. AD to control for the impact of degraded speech provided by the CIs.

##### 1.5.4 Relationship between neural tracking and implantation age

We performed a linear regression where age at which children received the first implant was the independent variable and the individual latency of the GFP-TRF first peak (GFP P1, we employed GFP to extract a more reliable peak latency ^23^) was the dependent variable.

##### 1.5.5 Relationship between neural tracking and speech comprehension

A point-by-point linear regression model was run to predict TRF values from behavioral accuracy of comprehension questions, groups (categorial variable), the interaction between them, and children’s age, to account for its impact, across the whole-time window [0 – 600 ms] at each electrode. We computed the averaged r^2^ across electrodes for each time-point, and we constructed a null distribution of r^2^ with 1000 permutations by shuffling the TRF values to compare at each time point the actual averaged r^2^ effect to the null averaged r^2^ distribution. The empirical p-values obtained were corrected in time with FDR.

### 2. Results

#### 2.1 Cohen’s d

To report the magnitude of the effect, we computed the unbiased estimate of Cohen’s d; we acknowledge that the effect can be inflated since it is computed on the same sample. Its confidence interval was computed with a bootstrap method with 1000 permutations. When multiple time points and/or channels were significant, Cohen’s d was computed on the mean of the time window in which the significant effect emerged and across the significant channels.

#### 2.2. The existence of TRF with respect to the null-TRF across all electrodes

Within each group, a cluster-based permutation test (same parameters as in the main analysis reported above) was performed to assess the difference between TRFs and the null-TRFs across all electrodes and time lags between 0 and 600 ms. In the HC group, four positive and two negative significant clusters emerged (p_clust_ < 0.05, see Figure S1A). In the CI group, two positive and two negative significant clusters emerged (all p_clust_ < 0.05, Figure S1B). In the HC-v group, three positive and two negative significant clusters emerged (p_clust_ < 0.05, see Figure S1C). Finally, when we tested the two subgroups of CI, CD, and AD separately, significant differences between the TRF and the null-TRF emerged. Within both CD and AD group, one positive and one negative significant cluster emerged (p_clust_ < 0.05, Figure S1D).

**Figure S1.**
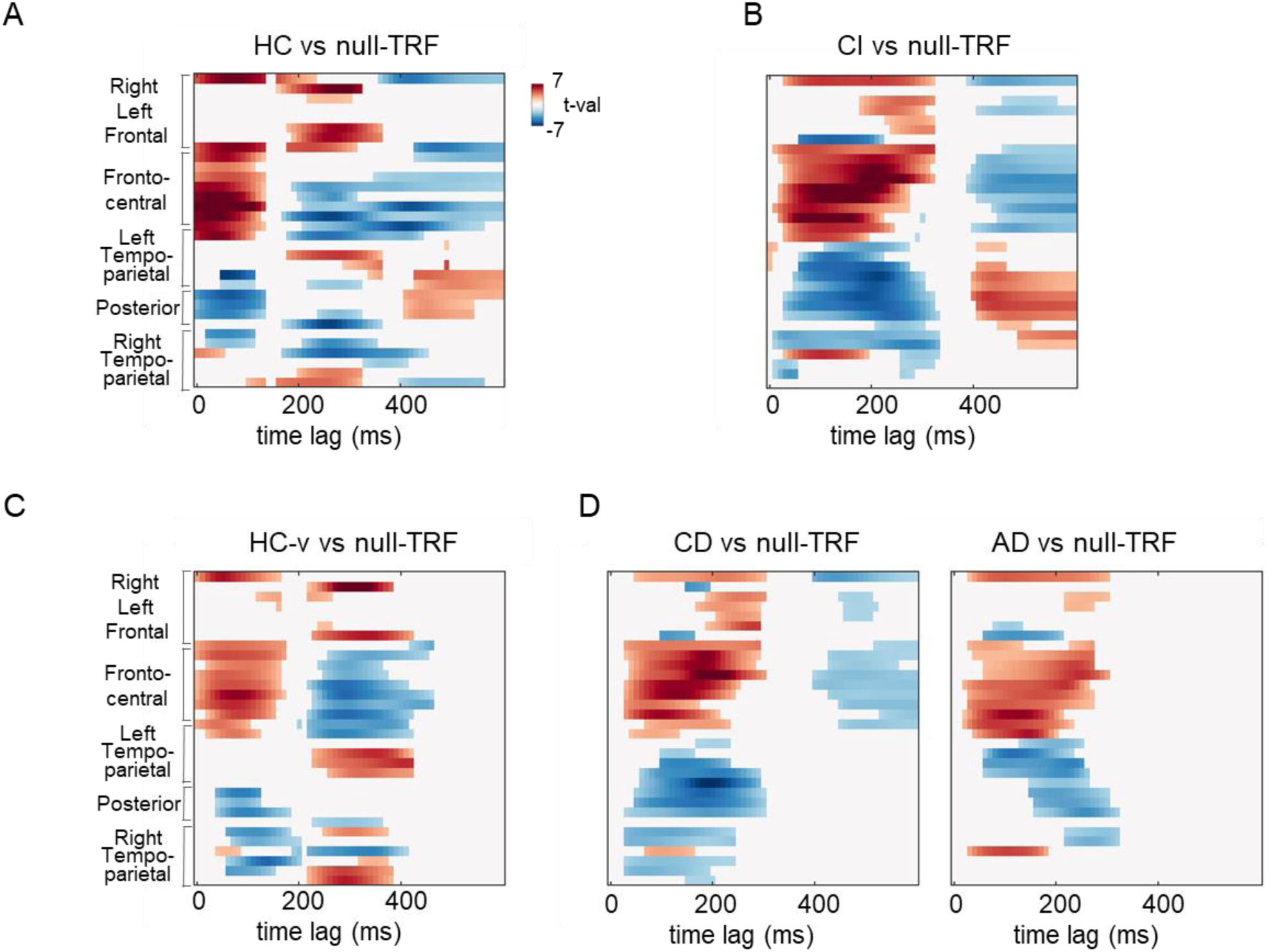
Existence of neural tracking of speech in all groups. Plots representing the results of cluster-based permutation tests contrasting the TRF and the null-TRF for HC, CI, HC-v, and CD and AD subgroups (A, B, C, and D panels respectively).

**Figure S2.**
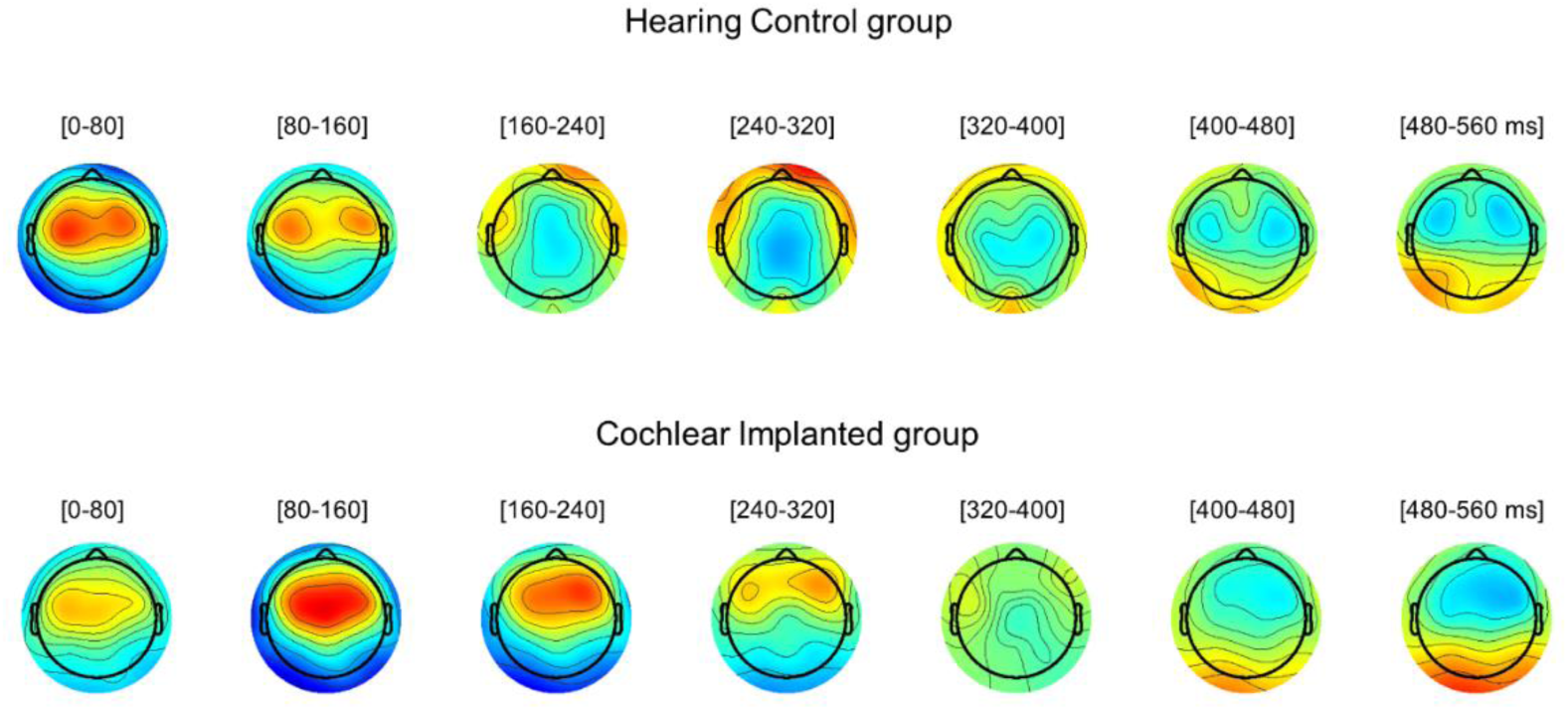
The spatiotemporal dynamics of HC’s and CI’s TRFs. Topography showing the TRFs across time (successive time window of 80 ms), separately for hearing control (HC) and cochlear implanted (CI) groups.

#### 2.3 Frontocentral TRF in CD, AD and HC-v

For both CD, AD, and HC-v, the frontocentral cluster TRF was significantly different from the null-TRF, highlighting a clear P1 (p_FDR_ < 0.05; see Figure S3).

**Figure S3.**
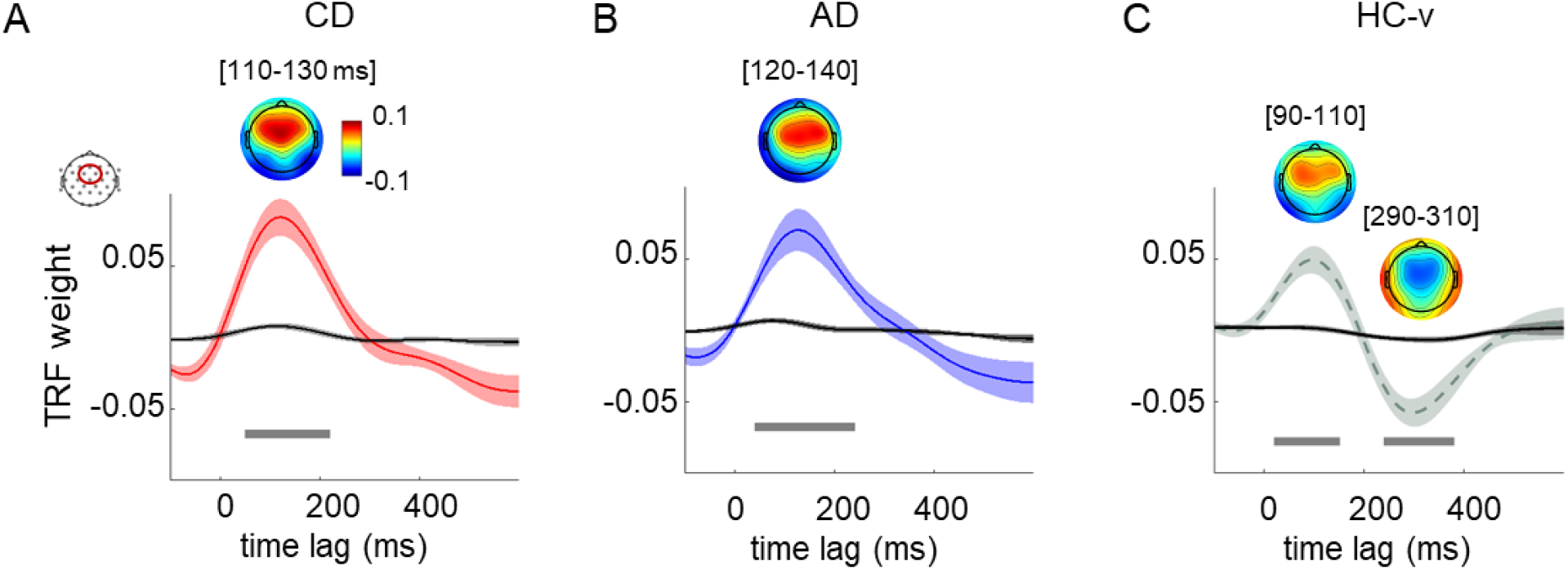
Frontocentral TRF in CD, AD, and HC-v groups. Grand average TRFs measured at frontocentral electrodes (Cz, Fz, FC1, and FC2) between -100 and 600 ms time lags are shown for CD, AD, and HC-v groups (A, B, C panel, respectively). Grand average null-TRFs computed for each group is plotted in grey. Shaded areas represent SE of the mean. Grey horizontal bars indicate time lags (between 0 and 600 ms) at which TRFs differed significantly from the null-TRF (running t-tests, FDR corrected p_FDR_ < 0.05).

#### 2.4 Test each CI group vs HC

Cluster-based permutation tests revealed that both CD and AD are significantly different from HC (one positive and one negative cluster, p_clust_<0.05).

#### 2.5 Relationship between GFP-TRF P1 latency and age at cochlear implantation

Given the high variability in the neural tracking latency (GFP-TRF P1) of the earliest implanted children, we explored whether a piecewise-linear regression model can significantly explain the data using the adaptive regression splines toolbox ^24^. We defined the best number of linear basis functions for our model using the pruning procedures. Thus, we tested the final model with the identified number of basis functions (i.e., 2, including the intercept). See Figure S4 for the results.

Moreover, we assessed whether some other clinical characteristics could explain the variance in TRF latency of children implanted before 21 months of age. We ran three separate simple linear regressions within this subgroup of CI participants who were implanted before 21 months of age to explore the impact on the P1 latency of chronological age, experience with the implant, and age at which hearing aids were provided before implantation, respectively. None of these factors explain the variance of the P1 latency in the earliest implanted children (all p > 0.05).

**Figure S4.**
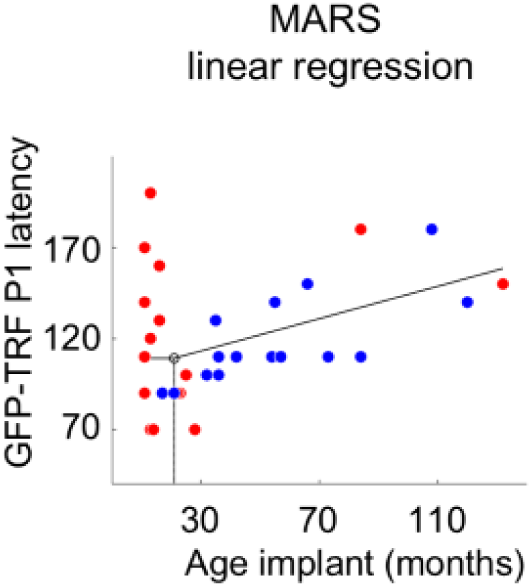
Relationship between neural tracking latency and implantation age. (A) The plot shows the piecewise-linear regression revealing that the positive relationship between implantation age and the latency of the neural tracking emerged from 21 months. Moreover, the color code (CD children are represented in red, while AD, in blue) highlights that most children implanted earlier than 21 months were diagnosed with bilateral profound congenital deafness.

#### 2.6 CI TRF temporally realigned

To account for the delay in the CI group, we realigned their first peak (P1) to the first peak of the HC group. The shift amount was 60 ms, equal to the difference between the mean of HC’s first peak (60 ms) and CI’s first peak (120 ms) computed on the frontocentral TRF cluster.

#### 2.7 Linear regression models (at all sensors) to predict the point-by-point TRF

We run a series of linear regression models (at all sensors) to predict the point-by-point TRF values from response accuracy and groups, their interaction, and age to account for its impact: TRF(c,t) = Accuracy + Group + Age + Accuracy × Group. Note that the CI’s TRF are shifted in time to account for the delay emerging at short timescale. As shown in the manuscript Figure 4D, the model emerged to be significant across electrodes between 130 and 300 ms. Figure S5 below shows where and when the single coefficients are significant. Clearly the relevant time window is around 200 ms (p < 0.05). Moreover, to explore the impact of accuracy within each group, we run the model TRF(c,t) = Accuracy + Age separately for HC and CI groups (see Figure S5B and C).

**Figure S5.**
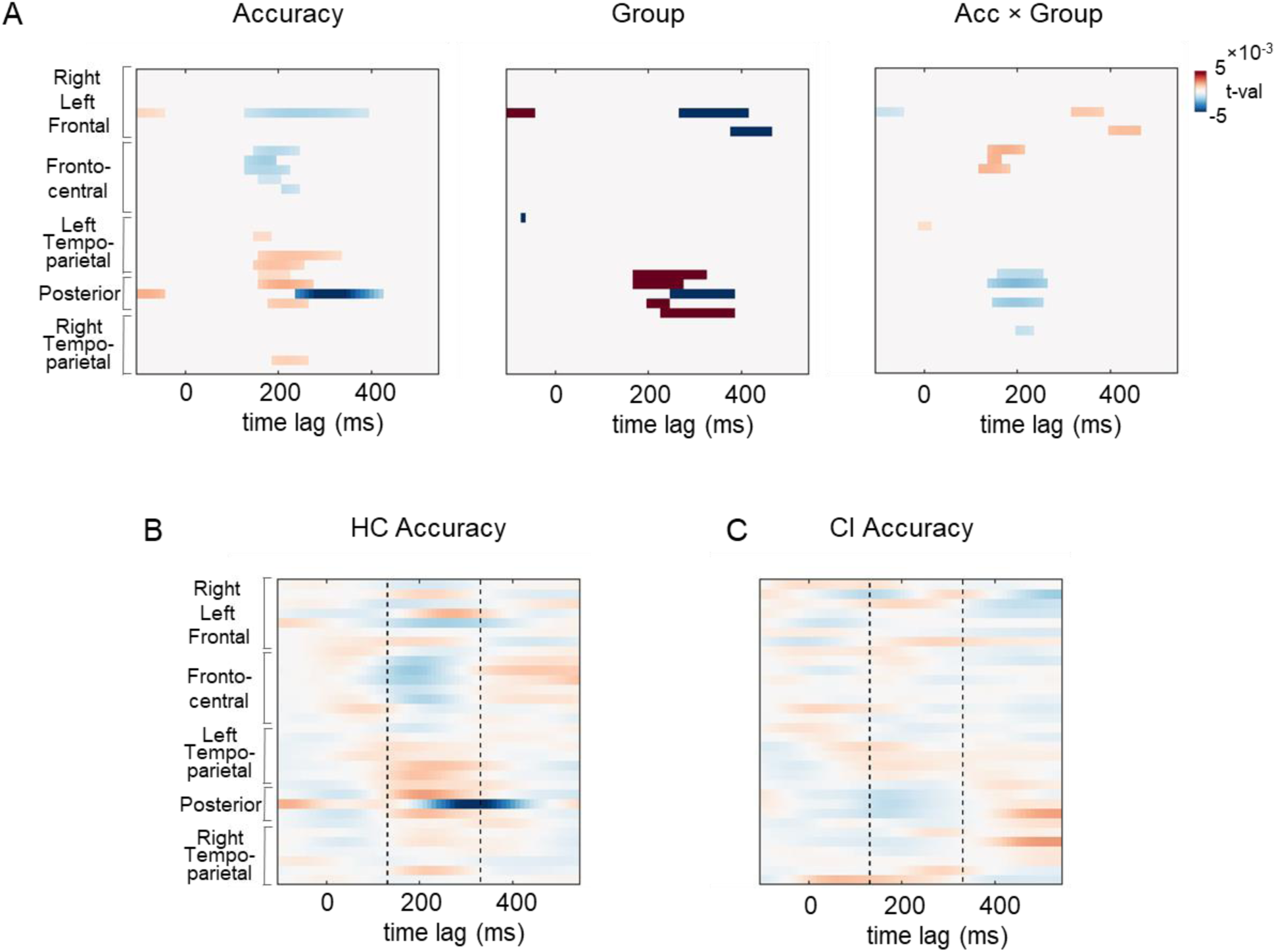
Channels by time coefficients of regression models. (A) The plots show the coefficient values with p < 0.05 for Accuracy, Group and their interaction obtained in the regression model in which the point-by-point TRF values were predicted by behavioral accuracy, group, the interaction between them, and age, introduced only to account for its impact. (B) and (D) panels shows the accuracy coefficients obtained by a linear regression model run separately within each group HC and CI, respectively. The dashed lines highlight the time window of interested from 130 to 330 ms.

#### 2.8 Results without cleaning of the artifacts

Comparable results emerged even without the artifact-cleaning procedure.

When we tested the existence of the auditory response function (between 0 and 600 ms), the frontocentral cluster TRF emerged to be significantly different from the null-TRF at two time windows between 20 and 260 ms and between 400 and 600 ms (p_FDR_ < 0.05).

Cluster-based permutation test between CI and HC groups performed at the whole brain level (across all channels) and comprising the TRF at every time lag between 0 to 600 ms revealed a significant difference between CI and HC groups (all p_clust_ < 0.05).

No difference emerged between CD and AD (no clusters were found at p_clust_ < 0.05).

**Table S1.**
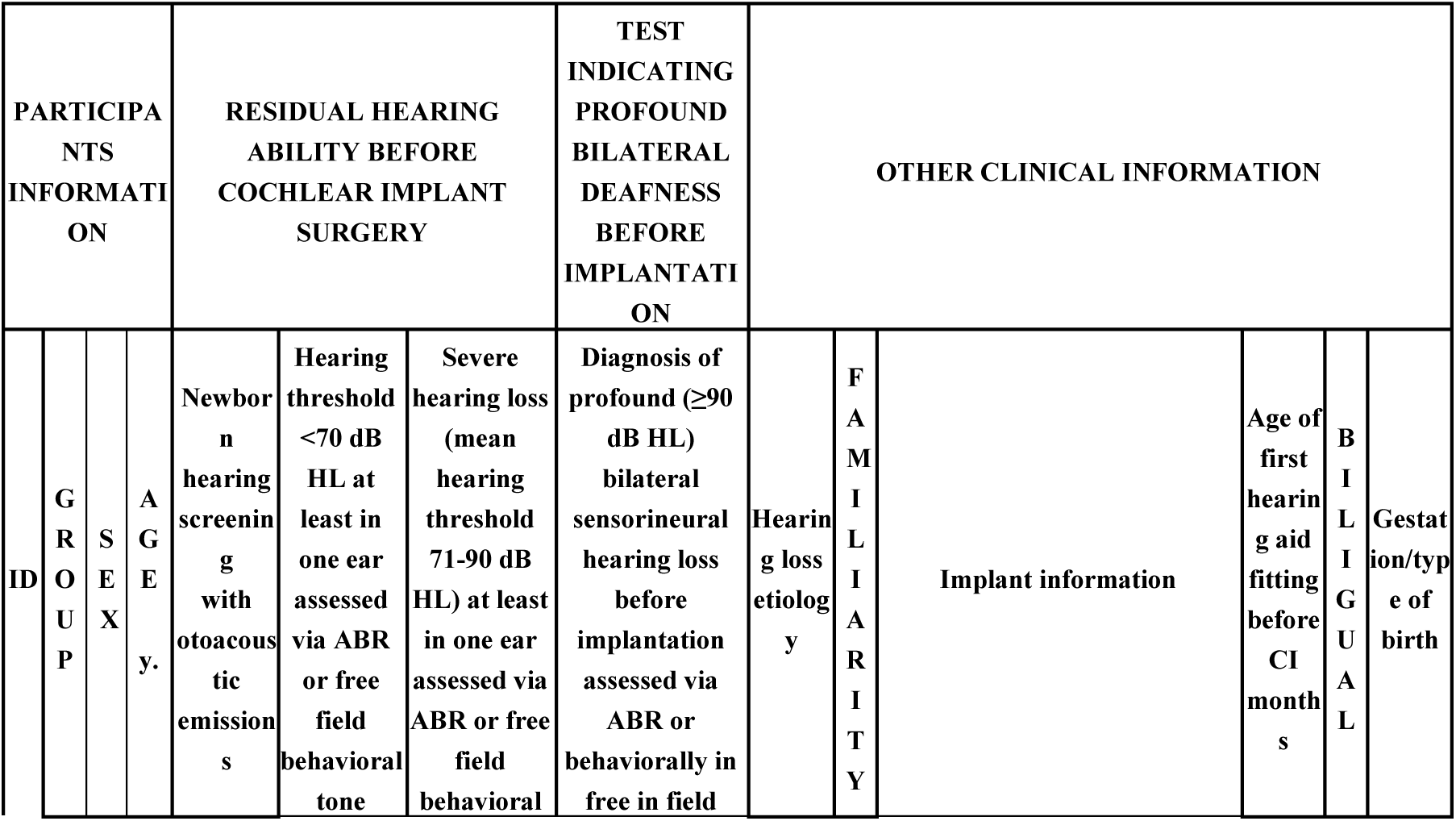
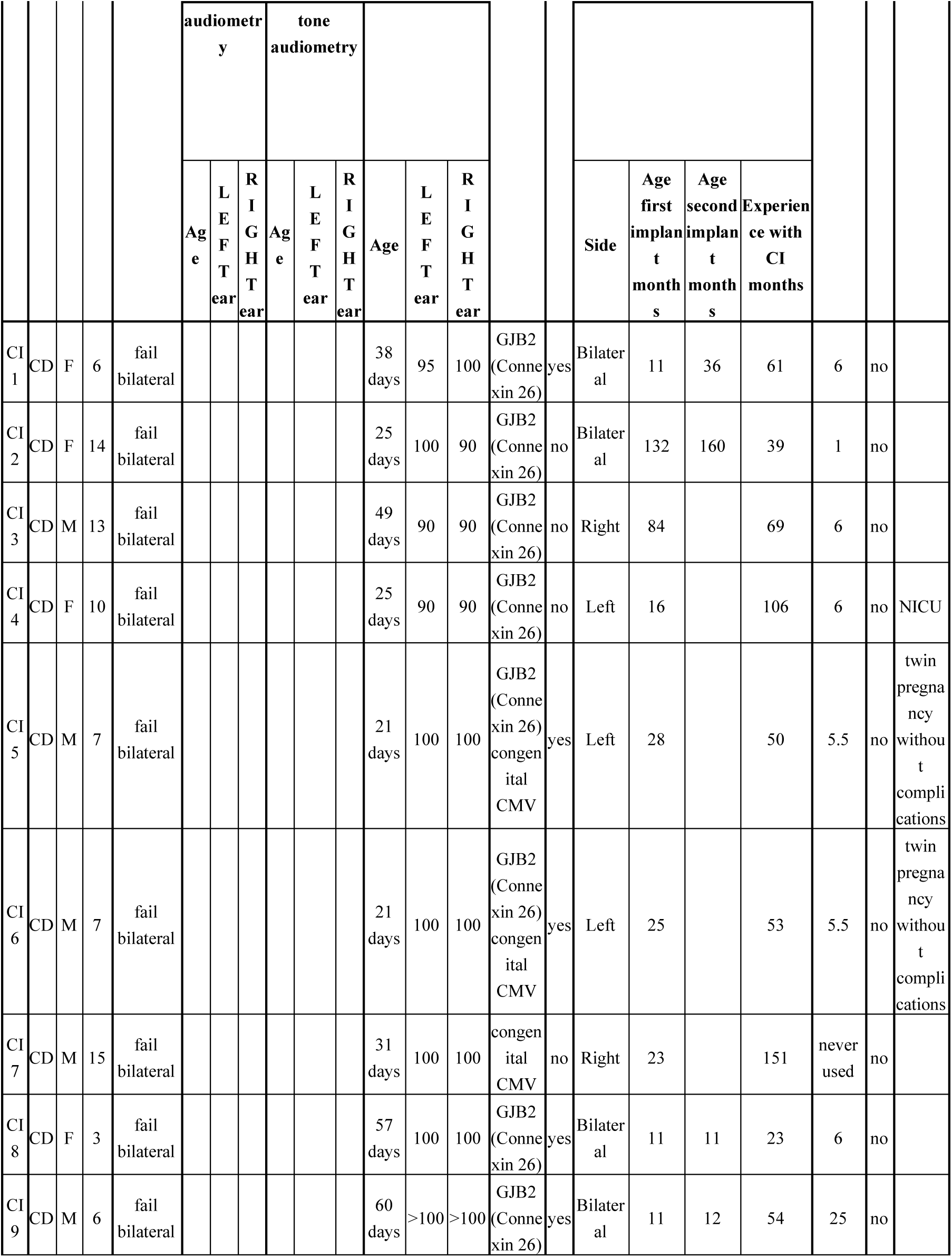
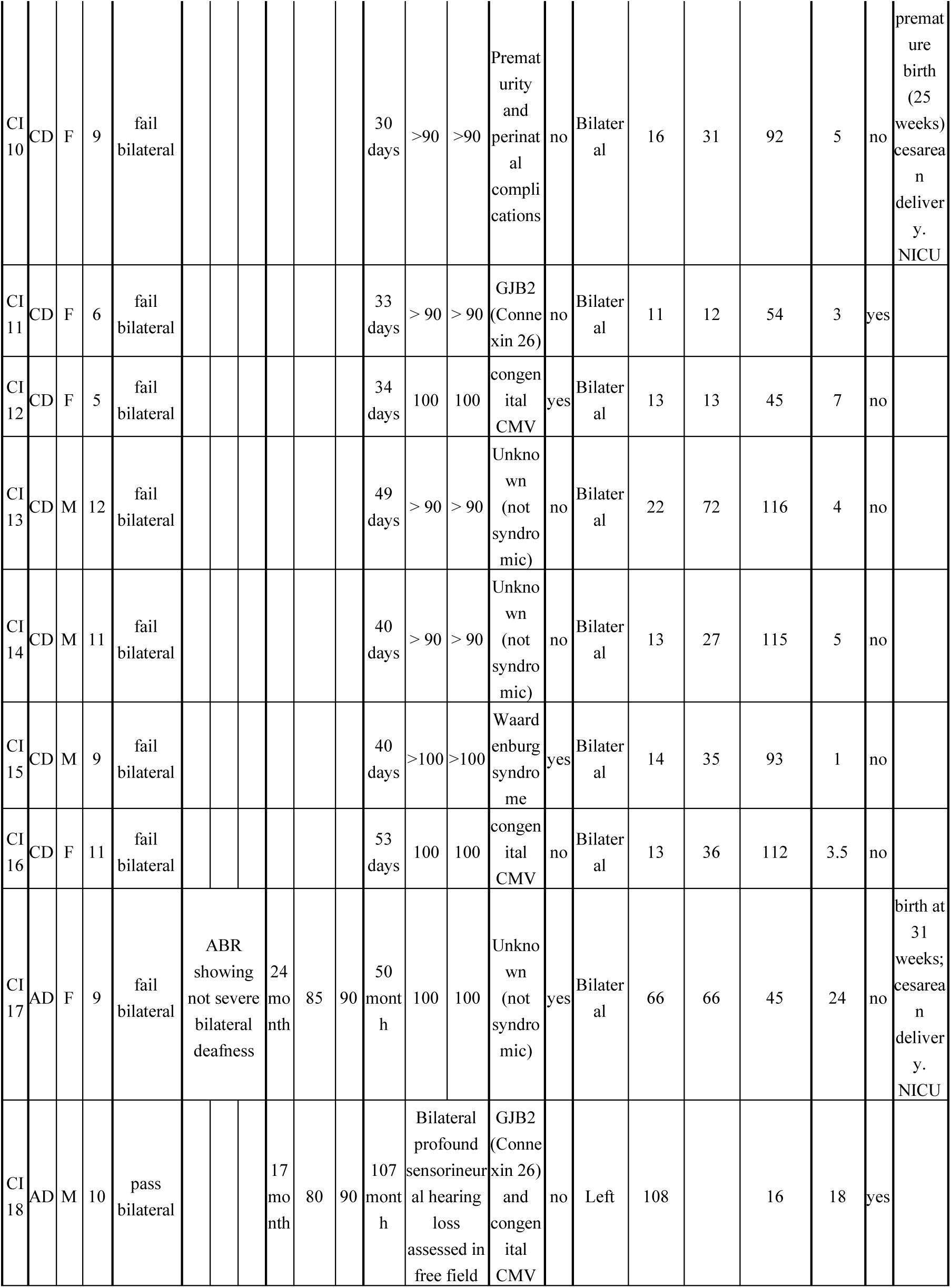
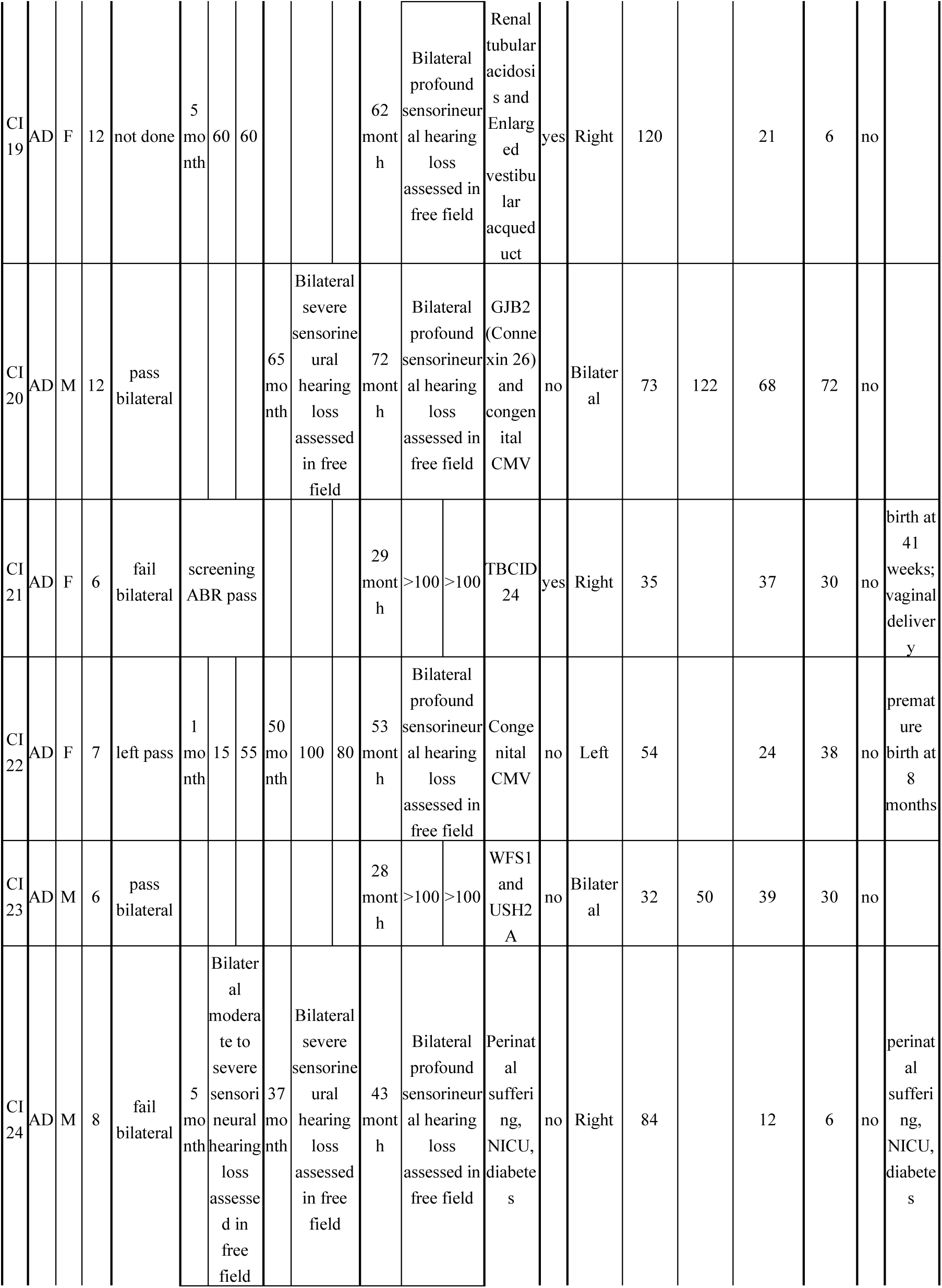
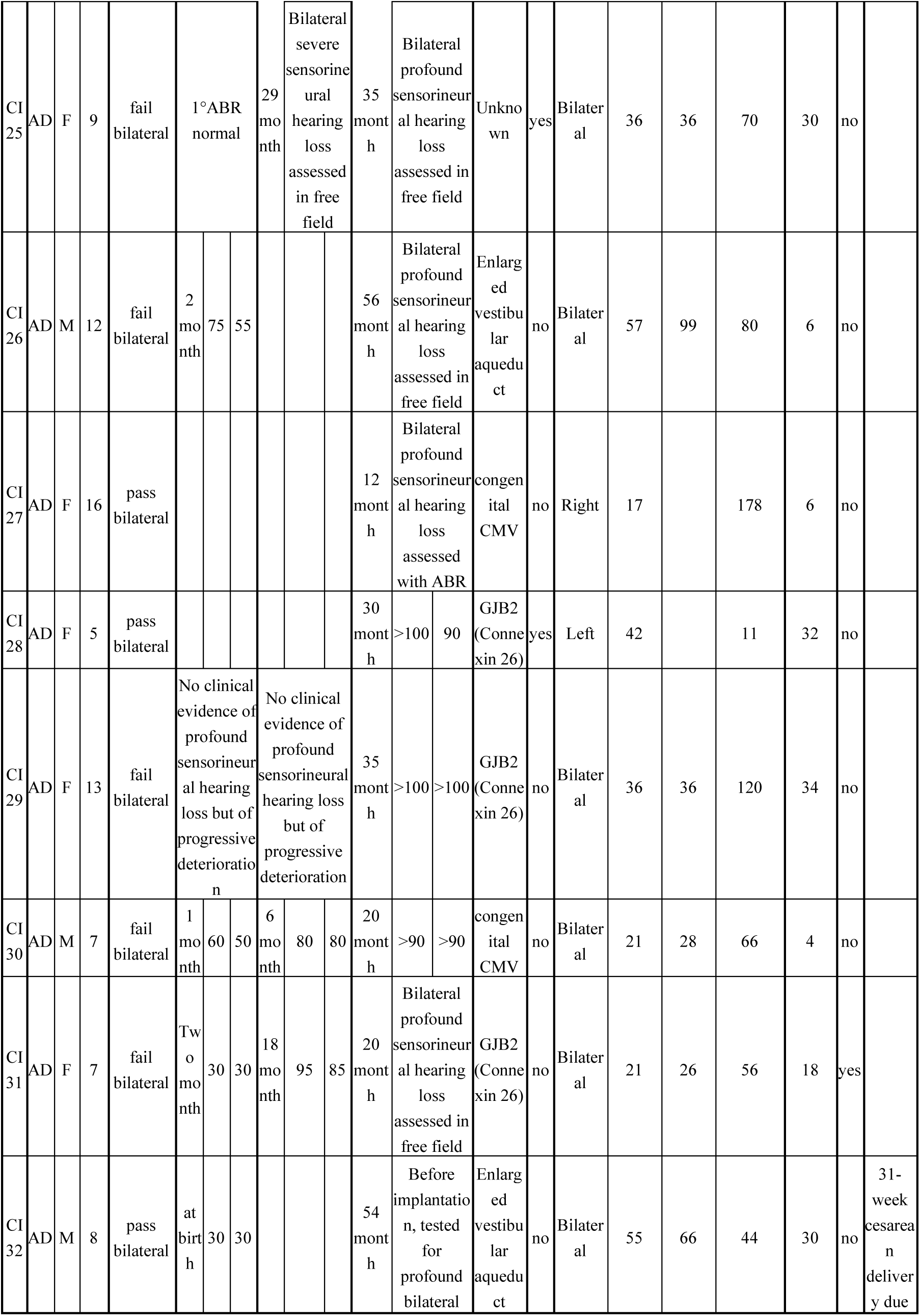
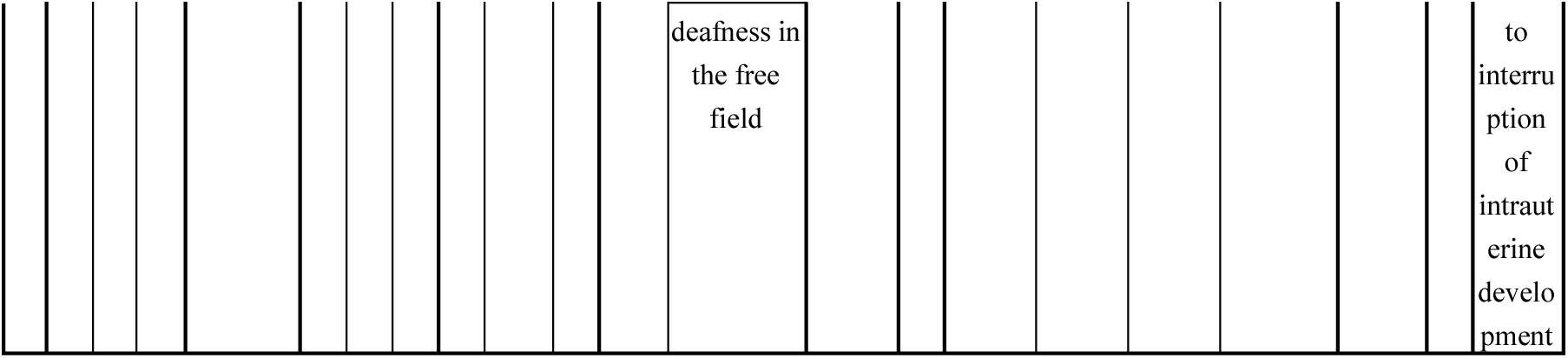
Table Characteristics of CI participants.

Characteristics of each CI participant included in the final sample. Degrees of hearing loss were defined following the American Speech-Language-Hearing Association (https://www.asha.org/public/hearing/degree-of-hearing-loss/, see also ^25^).

## References

1. Shannon R V, Zeng FG, Kamath V, Wygonski J, Ekelid M. Speech Recognition with Primarily Temporal Cues. Science (1979). 1995;270(5234):303–304. doi:10.1126/SCIENCE.270.5234.303

2. Drullman R, Festen JM, Plomp R. Effect of temporal envelope smearing on speech reception. J Acoust Soc Am. 1994;95(2):1053–1064. doi:10.1121/1.408467

3. Drullman R, Festen JM, Plomp R. Effect of reducing slow temporal modulations on speech reception. J Acoust Soc Am. 1994;95(5):2670–2680. doi:10.1121/1.409836

4. Kalashnikova M, Peter V, Di Liberto GM, Lalor EC, Burnham D. Infant-directed speech facilitates seven-month-old infants’ cortical tracking of speech. Sci Rep. 2018;8(1):1–8. doi:10.1038/s41598-018-32150-6

5. Jessen S, Fiedler L, Münte TF, Obleser J. Quantifying the individual auditory and visual brain response in 7-month-old infants watching a brief cartoon movie. Neuroimage. 2019;202(April):116060. doi:10.1016/j.neuroimage.2019.116060

6. Ortiz Barajas MC, Guevara R, Gervain J. The origins and development of speech envelope tracking during the first months of life. Dev Cogn Neurosci. 2021;48. doi:10.1016/j.dcn.2021.100915

7. Attaheri A, Choisdealbha ÁN, Di Liberto GM, et al. Delta- and theta-band cortical tracking and phase-amplitude coupling to sung speech by infants. Neuroimage. 2022;247(November 2021):118698. doi:10.1016/j.neuroimage.2021.118698

8. Reh RK, Dias BG, Nelson CA, et al. Critical period regulation acrossmultiple timescales. Proc Natl Acad Sci U S A. 2020;117(38):23242–23251. doi:10.1073/pnas.1820836117

9. Werker JF, Hensch TK. Critical periods in speech perception: New directions. Annu Rev Psychol. 2015;66:173–196. doi:10.1146/annurev-psych-010814-015104

10. Bottari D, Berto M. Three factors to characterize plastic potential transitions in the visual system. Neurosci Biobehav Rev. 2021;126(November 2020):444–446. doi:10.1016/j.neubiorev.2021.03.035

11. Hensch TK. Critical period plasticity in local cortical circuits. Nat Rev Neurosci. 2005;6(11):877–888. doi:10.1038/nrn1787

12. Winn MB, Nelson PB. Cochlear Implants. Oxford Research Encyclopedia of Linguistics. Published online August 31, 2021. doi:10.1093/ACREFORE/9780199384655.013.893

13. Pavani F, Bottari D. Neuroplasticity following cochlear implants. Handb Clin Neurol. 2022;187:89–108. doi:10.1016/B978-0-12-823493-8.00016-X

14. Gates GA, Daly K, Dichtel WJ, et al. Cochlear Implants in Adults and Children. JAMA. 1995;274(24):1955–1961. doi:10.1001/JAMA.1995.03530240065043

15. Clark G. Cochlear Implants: Fundamentals and Applications. Springer New York.; 2003.

16. Tamati TN, Pisoni DB, Moberly AC. Speech and language outcomes in adults and children with cochlear implants. Annual Review of Linguistics. 2022; 8(1), 299–319.

17. Houston DM, Stewart J, Moberly A, Hollich G, Miyamoto RT. Word learning in deaf children with cochlear implants: Effects of early auditory experience. Dev Sci. 2012;15(3):448–461. doi:10.1111/j.1467-7687.2012.01140.x

18. Niparko JK, Tobey EA, Thal DJ, et al. Spoken language development in children following cochlear implantation. JAMA. 2010;303(15):1498–1506. doi:10.1001/jama.2010.451

19. Sharma SD, Cushing SL, Papsin BC, Gordon KA. Hearing and speech benefits of cochlear implantation in children: A review of the literature. International journal of pediatric otorhinolaryngology. 2020;133, 109984.

20. Kral A, Dorman MF, Wilson BS. Neuronal development of hearing and language: cochlear implants and critical periods. Annual review of neuroscience. 2019; 42(1), 47–65.

21. Sharma A, Dorman M, Spahr A, Todd NW. Early cochlear implantation in children allows normal development of central auditory pathways. *Annals of Otology*, Rhinology and Laryngology. 2002;111(5 II):38–41. doi:10.1177/00034894021110s508

22. Sharma A, Dorman M, Spahr A. A sensitive period for the development of the central auditory system in children with cochlear implants: Implications for age of implantation. Ear Hear. 2002;23(6):532–539. doi:10.1097/00003446-200212000-00004

23. Sharma A, Dorman MF, Kral A. The influence of a sensitive period on central auditory development in children with unilateral and bilateral cochlear implants. Hear Res. 2005;203(1-2):134–143. doi:10.1016/j.heares.2004.12.010

24. Sharma A, Campbell J, Cardon G. Developmental and cross-modal plasticity in deafness: evidence from the P1 and N1 event related potentials in cochlear implanted children. Int J Psychophysiol. 2015;95(2):135–144. doi:10.1016/J.IJPSYCHO.2014.04.007

25. Eggermont JJ, Ponton CW. Auditory-evoked potential studies of cortical maturation in normal hearing and implanted children: Correlations with changes in structure and speech perception. Acta Otolaryngol. 2003;123(2):249–252. doi:10.1080/0036554021000028098

26. Delorme A, Makeig S. EEGLAB: An open source toolbox for analysis of single-trial EEG dynamics including independent component analysis. J Neurosci Methods. 2004;134(1):9–21. doi:10.1016/j.jneumeth.2003.10.009

27. Stropahl M, Bauer AKR, Debener S, Bleichner MG. Source-Modeling auditory processes of EEG data using EEGLAB and brainstorm. Front Neurosci. 2018;12(MAY):348523. doi:10.3389/FNINS.2018.00309/BIBTEX

28. Bottari D, Bednaya E, Dormal G, et al. EEG frequency-tagging demonstrates increased left hemispheric involvement and crossmodal plasticity for face processing in congenitally deaf signers. Neuroimage. 2020;223(December 2019):117315. doi:10.1016/j.neuroimage.2020.117315

29. Deprez H, Gransier R, Hofmann M, van Wieringen A, Wouters J, Moonen M. Characterization of cochlear implant artifacts in electrically evoked auditory steady-state responses. Biomed Signal Process Control. 2017;31:127–138. doi:10.1016/j.bspc.2016.07.013

30. Somers B, Verschueren E, Francart T. Neural tracking of the speech envelope in cochlear implant users. J Neural Eng. 2019;16(1):1–23. doi:10.1088/1741-2552/aae6b9

31. Mirkovic B, Debener S, Jaeger M, De Vos M. Decoding the attended speech stream with multi-channel EEG: Implications for online, daily-life applications. J Neural Eng. 2015;12(4). doi:10.1088/1741-2560/12/4/046007

32. O’Sullivan JA, Power AJ, Mesgarani N, et al. Attentional Selection in a Cocktail Party Environment Can Be Decoded from Single-Trial EEG. Cerebral Cortex. 2015;25(7):1697–1706. doi:10.1093/cercor/bht355

33. Crosse MJ, Di Liberto GM, Bednar A, Lalor EC. The multivariate temporal response function (mTRF) toolbox: A MATLAB toolbox for relating neural signals to continuous stimuli. Front Hum Neurosci. 2016;10(NOV2016):1–14. doi:10.3389/fnhum.2016.00604

34. Combrisson E, Jerbi K. Exceeding chance level by chance: The caveat of theoretical chance levels in brain signal classification and statistical assessment of decoding accuracy. J Neurosci Methods. 2015;250:126–136. doi:10.1016/j.jneumeth.2015.01.010

35. Crosse MJ, Zuk NJ, Di Liberto GM, Nidiffer AR, Molholm S, Lalor EC. Linear Modeling of Neurophysiological Responses to Speech and Other Continuous Stimuli: Methodological Considerations for Applied Research. Front Neurosci. 2021;15. doi:10.3389/fnins.2021.705621

36. APA Yu AB, Hairston WD. Open EEG Phantom. Published January 4, 2021. Accessed July 31, 2024. 10.17605/OSF.IO/QRKA2

37. Paul BT, Uzelac M, Chan E, Dimitrijevic A. Poor early cortical differentiation of speech predicts perceptual difficulties of severely hearing-impaired listeners in multi-talker environments. Sci Rep. 2020;10(1):1–12. doi:10.1038/s41598-020-63103-7

38. Steinschneider M, Liégeois-Chauvel C, Brugge JF. Auditory Evoked Potentials and Their Utility in the Assessment of Complex Sound Processing. In: Springer, Boston M, ed. The Auditory Cortex. ; 2011:535–559. doi:10.1007/978-1-4419-0074-6

39. Benjamini Y, Yekutieli D. The Control of the False Discovery Rate in Multiple Testing under Dependency. Vol 29.; 2001.

40. Maris E, Oostenveld R. Nonparametric statistical testing of EEG- and MEG-data. J Neurosci Methods. 2007;164(1):177–190. doi:10.1016/j.jneumeth.2007.03.024

41. Fiedler L, Wöstmann M, Herbst SK, Obleser J. Late cortical tracking of ignored speech facilitates neural selectivity in acoustically challenging conditions. Neuroimage. 2019;186:33–42. doi:10.1016/J.NEUROIMAGE.2018.10.057

42. Barriga-Paulino CI, Rodríguez-Martínez EI, Arjona A, Morales M, Gómez CM. Developmental trajectories of event related potentials related to working memory. Neuropsychologia. 2017;95:215–226. doi:10.1016/j.neuropsychologia.2016.12.026

43. Watanabe T, Rees G, Masuda N. Atypical intrinsic neural timescale in autism. Elife. 2019;8. doi:10.7554/ELIFE.42256

44. Watanabe T. Causal roles of prefrontal cortex during spontaneous perceptual switching are determined by brain state dynamics. Elife. 2021;10. doi:10.7554/ELIFE.69079

45. Lehmann D, Skrandies W. Reference-free identification of components of checkerboard-evoked multichannel potential fields. Electroencephalogr Clin Neurophysiol. 1980;48(6):609–621. doi:10.1016/0013-4694(80)90419-8

46. Michel CM. Electrical Neuroimaging. Cambridge University Press; 2009.

47. Vanthrnhout J, Decruy L, Wouters J, Simon JZ, Francart T. Vanthrnhout et al., 2017_Speech Intelligibility Predicted from Neural Entrainment of the Speech Envelope.pdf. Published online 2018:181–191.

48. Pérez-Navarro J, Klimovich-Gray A, Lizarazu M, Piazza G, Molinaro N, Lallier M. The contribution of early language exposure to the cortical tracking of speech. BioRxiv. Published online 2023. doi:10.1101/2023.09.14.557701

49. Ortiz-Barajas MC, Guevara R, Gervain J. Neural oscillations and speech processing at birth. iScience. 2023;26(11). doi:10.1016/j.isci.2023.108187

50. Gillis M, Decruy L, Vanthornhout J, Francart T. Hearing loss is associated with delayed neural responses to continuous speech. European Journal of Neuroscience. 2022; 55(6), 1671–1690.

51. Billings CJ, Bennett KO, Molis, MR, Leek MR. Cortical encoding of signals in noise: effects of stimulus type and recording paradigm. Ear Hear. 2011;32(1):53–60.

52. Ding N, Simon JZ. Adaptive temporal encoding leads to a background-insensitive cortical representation of speech. Journal of Neuroscience. 2013;33(13):5728–5735. doi:10.1523/JNEUROSCI.5297-12.2013

53. Gustafson SJ, Billings CJ, Hornsby BWY, Key AP. Effect of competing noise on cortical auditory evoked potentials elicited by speech sounds in 7-to 25-year-old listeners. Hear Res. 2019;373:103–112. doi:10.1016/j.heares.2019.01.004

54. Yasmin S, Irsik VC, Johnsrude IS, Herrmann B. The effects of speech masking on neural tracking of acoustic and semantic features of natural speech. Neuropsychologia. 2023;186:108584. doi:10.1016/J.NEUROPSYCHOLOGIA.2023.108584

55. Kral A, Sharma A. Developmental neuroplasticity after cochlear implantation. Trends Neurosci. 2012;35(2):111–122. doi:10.1016/J.TINS.2011.09.004

56. Sharma A, Campbell J. A sensitive period for cochlear implantation in deaf children. J Matern Fetal Neonatal Med. 2011;24(1):151. doi:10.3109/14767058.2011.607614

57. Sharma A, Gilley PM, Dorman MF, Baldwin R. Deprivation-induced cortical reorganization in children with cochlear implants. Int J Audiol. 2007;46(9):494–499. doi:10.1080/14992020701524836

58. Etard O, Reichenbach T. Neural Speech Tracking in the Theta and in the Delta Frequency Band Differentially Encode Clarity and Comprehension of Speech in Noise. The Journal of Neuroscience. 2019;39(29):5750. doi:10.1523/JNEUROSCI.1828-18.2019

59. Chen YP, Schmidt F, Keitel A, Rösch S, Hauswald A, Weisz N. Speech intelligibility changes the temporal evolution of neural speech tracking. NeuroImage. 2023; 268, 119894.

60. Lieu JEC, Kenna M, Anne S, Davidson L. Hearing Loss in Children: A Review. JAMA. 2020;324(21):2195–2205. doi:10.1001/JAMA.2020.17647

61. Mariani B, Nicoletti G, Barzon G, et al. Prenatal experience with language shapes the brain. Sci Adv. 2023;9(47). doi:10.1126/SCIADV.ADJ3524

## References

1. Fiedler L, Wöstmann M, Herbst SK, Obleser J. Late cortical tracking of ignored speech facilitates neural selectivity in acoustically challenging conditions. Neuroimage. 2019;186:33–42. doi:10.1016/J.NEUROIMAGE.2018.10.057

2. Wang C, Zhang Q. Word frequency effect in written production: Evidence from ERPs and neural oscillations. Psychophysiology. 2021;58(5):e13775. doi:10.1111/PSYP.13775

3. 3. Berrettini S, BAggiAni A, Burdo S, et al. Analysis of the Impact of Professional Involvement in Evidence Generation for the HTA Process, Subproject “Cochlear Implants”: Methodology, Results and Recommendations Analisi Dell’impatto Del Coinvolgimento Dei Professionisti Nella Produzione Dell’evidenza per i Processi Di HTA, Sottoprogetto “Impianti Cocleari”: Metodologia, Risultati, Raccomandazioni. Vol 31.; 2011. https://sites.google.com/site/impianticoclear-

4. Delorme A, Sejnowski T, Makeig S. Enhanced detection of artifacts in EEG data using higher-order statistics and independent component analysis. Neuroimage. 2007;34(4):1443–1449. doi:10.1016/J.NEUROIMAGE.2006.11.004

5. Bell AJ, Sejnowski TJ. An information-maximization approach to blind separation and blind deconvolution. Neural Comput. 1995;7(6):1129–1159. doi:10.1162/NECO.1995.7.6.1129

6. Jung TP, Makeig S, Humphries C, et al. Removing electroencephalographic artifacts by blind source separation. Psychophysiology. 2000;37(2):163–178. doi:10.1111/1469-8986.3720163

7. Jung TP, Makeig S, Westerfield M, Townsend J, Courchesne E, Sejnowski TJ. Removal of eye activity artifacts from visual event-related potentials in normal and clinical subjects. Clinical Neurophysiology. 2000;111(10):1745–1758. doi:10.1016/S1388-2457(00)00386-2

8. Stropahl M, Bauer AKR, Debener S, Bleichner MG. Source-Modeling auditory processes of EEG data using EEGLAB and brainstorm. Front Neurosci. 2018;12(MAY):348523. doi:10.3389/FNINS.2018.00309/BIBTEX

9. Bottari D, Bednaya E, Dormal G, et al. EEG frequency-tagging demonstrates increased left hemispheric involvement and crossmodal plasticity for face processing in congenitally deaf signers. Neuroimage. 2020;223(December 2019):117315. doi:10.1016/j.neuroimage.2020.117315

10. Viola FC, Thorne J, Edmonds B, Schneider T, Eichele T, Debener S. Semi-automatic identification of independent components representing EEG artifact. Clinical Neurophysiology. 2009;120(5):868–877. doi:10.1016/j.clinph.2009.01.015

11. Paul BT, Uzelac M, Chan E, Dimitrijevic A. Poor early cortical differentiation of speech predicts perceptual difficulties of severely hearing-impaired listeners in multi-talker environments. Sci Rep. 2020;10(1):1–12. doi:10.1038/s41598-020-63103-7

12. Crosse MJ, Zuk NJ, Di Liberto GM, Nidiffer AR, Molholm S, Lalor EC. Linear Modeling of Neurophysiological Responses to Speech and Other Continuous Stimuli: Methodological Considerations for Applied Research. Front Neurosci. 2021;15. doi:10.3389/fnins.2021.705621

13. Crosse MJ, Di Liberto GM, Bednar A, Lalor EC. The multivariate temporal response function (mTRF) toolbox: A MATLAB toolbox for relating neural signals to continuous stimuli. Front Hum Neurosci. 2016;10(NOV 2016):1–14. doi:10.3389/fnhum.2016.00604

14. Lalor EC, Power AJ, Reilly RB, Foxe JJ. Resolving precise temporal processing properties of the auditory system using continuous stimuli. J Neurophysiol. 2009;102(1):349–359. doi:10.1152/jn.90896.2008

15. APA Yu AB, Hairston WD. Open EEG Phantom. Published January 4, 2021. Accessed July 31, 2024. 10.17605/OSF.IO/QRKA2

16. Steinschneider M, Liégeois-Chauvel C, Brugge JF. Auditory Evoked Potentials and Their Utility in the Assessment of Complex Sound Processing. In: Springer, Boston M, ed. The Auditory Cortex. ; 2011:535–559. doi:10.1007/978-1-4419-0074-6

17. Jessen S, Fiedler L, Münte TF, Obleser J. Quantifying the individual auditory and visual brain response in 7-month-old infants watching a brief cartoon movie. Neuroimage. 2019;202(April):116060. doi:10.1016/j.neuroimage.2019.116060

18. Benjamini Y, Yekutieli D. The Control of the False Discovery Rate in Multiple Testing under Dependency. Vol 29.; 2001.

19. Maris E, Oostenveld R. Nonparametric statistical testing of EEG- and MEG-data. J Neurosci Methods. 2007;164(1):177–190. doi:10.1016/j.jneumeth.2007.03.024

20. Oostenveld R, Fries P, Maris E, Schoffelen JM. FieldTrip: Open source software for advanced analysis of MEG, EEG, and invasive electrophysiological data. Comput Intell Neurosci. 2011;2011. doi:10.1155/2011/156869

21. Lorenzi C, Berthommier F, Demany L. Discrimination of amplitude-modulation phase spectrum. J Acoust Soc Am. 1999;105(5):2987–2990. doi:10.1121/1.426911

22. Strickland EA, Viemeister NF. Cues for discrimination of envelopes. J Acoust Soc Am. 1996;99(6):3638–3646. doi:10.1121/1.414962

23. Michel CM. Electrical Neuroimaging. Cambridge University Press; 2009.

24. Friedman JH. Multivariate adaptive regression splines. The annals of statistics. 1991;19(1):1–67.

25. Lieu JEC, Kenna M, Anne S, Davidson L. Hearing Loss in Children: A Review. JAMA. 2020;324(21):2195–2205. doi:10.1001/JAMA.2020.17647

